# A Sequential Assembly Mechanism for Stable Cdc13 Dimerization on Telomeric DNA

**DOI:** 10.64898/2026.09.01.748474

**Authors:** Po-Wei Chiu, Yu-Ting Lin, Pei-Ru Lai, Tzu-Yu Lee, Yu-Chen Cheng, Yan-Zhu Hsieh, Hung-Wen Li, Jing-Jer Lin

## Abstract

The telomere-binding protein Cdc13 specifically binds to single-stranded telomeric DNA, playing a critical role in telomere protection and length regulation. While extensive biochemical, molecular biological, and genetic studies have shown that Cdc13 can form dimers or oligomers in solution and bind telomeric DNA with high specificity, the dynamic mechanism of its loading onto telomeres is less well characterized. Combining two single-molecule methods, single-molecule fluorescence resonance energy transfer (smFRET) and colocalization single-molecule spectroscopy (CoSMoS), we demonstrate a kinetically favored sequential assembly pathway on a 12-nucleotide telomeric DNA segment involving an initial Cdc13 binding event followed by recruitment of a second Cdc13 molecule. While genetic studies suggest that monomeric Cdc13 binding alone is insufficient to maintain telomere length, our findings underscore the Cdc13 monomer’s regulatory importance in coordinating telomere synthesis and protection. This monomer-to-dimer transition provides a mechanistic basis for understanding the multifunctional roles of Cdc13 in telomere replication and protection.

**Key Points:**

- Two complementary single-molecule fluorescence methods, FRET and multi-color colocalization, are applied to characterize Cdc13 binding to telomeric DNA in real time.
- Monomeric Cdc13 binds sequentially to telomeric DNA, potentially leading to the formation of a relatively stable Cdc13 dimer-DNA complex.
- This sequential binding model may provide a mechanism for regulating Cdc13-telomere complex formation, contributing to telomere homeostasis.

## Introduction

Telomeres, located at the ends of eukaryotic chromosomes, are critical for cells to differentiate chromosomal ends from broken DNA ends, protect chromosomes from degradation by nucleases, and prevent end-to-end fusion of chromosomes. The functions of telomeres are executed by specific protein factors located on telomeres. Cell division cycle 13 (Cdc13) protein is the telomere-binding protein of yeast *Saccharomyces cerevisiae* (1,2). It binds specifically to single-stranded telomeric TG_1-3_ DNA (1–3). This specific binding protects telomeres from degradation and serves as a key regulatory platform for recruiting various proteins that execute different functions on telomeres (4). Cdc13 has multiple regulatory roles: it forms a CST complex with Stn1 and Ten1 to cap telomeres and negatively regulate telomere length (5,6); through interaction with Est1, a telomerase-associated protein, it stimulates telomerase activity (7–11); and by interacting with the catalytic subunit of DNA polymerase α, it mediates the C-strand synthesis of telomeres (8,12,13).

*S. cerevisiae CDC13* encodes a 924-amino-acid protein containing five functional domains: four oligonucleotide/oligosaccharide-binding (OB) folds and a recruitment domain (RD) (14,15). Each domain plays specific roles in telomere biology. Both the N-terminal OB1 and OB2 are involved in dimer formation (13,16,17). The OB1 domain also mediates binding to Pol1 (13), while OB2 participates in Stn1 binding (17). The OB3 contains a telomeric DNA binding region, and the RD interacts with Est1 to facilitate telomere extension by telomerase (4,18,19). Analysis of the *Kluyveromyces lactis* CST complex revealed that Cdc13 utilizes both the OB2 and OB4 domains to interact with Stn1 (15). Similar interactions likely occur between the OB2 and OB4 of *S. cerevisiae* Cdc13 and Stn1 (17,18). Interestingly, biochemical and structural analyses have demonstrated that Cdc13 forms dimers in solution (13,16,17), leading to the widely held assumption that Cdc13 binds directly to telomeres as a dimer. However, yeast cells harboring OB1-mutated Cdc13 exhibit short telomere length and temperature-sensitive growth (13), suggesting that while Cdc13 monomers can function to protect telomeres and support telomere extension, they are less effective than dimeric Cdc13. These findings may also indicate the presence of a Cdc13 monomer binding step during Cdc13 loading onto telomeres. Despite these insights, the detailed mechanistic process by which Cdc13 interacts with telomeric DNA remains poorly understood.

Given the important roles of Cdc13 in protecting telomere ends and maintaining telomere length, Cdc13 binding must be specific to telomeres with high affinity. Structural analysis revealed specific interactions between Cdc13 and telomeric DNA (19). The population-averaged biochemical studies determined specific and high-affinity binding of Cdc13 to telomeric DNA. However, direct measurement for the dynamic binding of Cdc13 to telomeres is lacking. Here we applied two complementary single-molecule techniques, fluorescence resonance energy transfer (smFRET) and colocalization single-molecule spectroscopy (CoSMoS), to directly characterize the binding kinetics and its oligomeric form. We found Cdc13 loads onto telomeric DNA as a monomer in a sequential manner to form a Cdc13 dimer-DNA complex. While the binding of monomeric Cdc13 binds to DNA is dynamic, the dimeric Cdc13-DNA structure is less susceptible to dissociation during 100 mM NaCl wash. This sequential binding of monomeric Cdc13 offers advantageous strategies for a sequence-specific and high-affinity association that allows Cdc13 to protect and regulate telomeres.

## Results

### Cdc13 binding to TG12 telomere DNA leads to two bound states in FRET

During most phases of the cell cycle in yeast, the size of single-stranded telomeres is reported to be ∼12-14 nucleotides (nt) long (20). Previous studies suggested that a 11-12 nt telomeric DNA is the minimum length required for Cdc13 binding (21). Thus, to study the binding of Cdc13 to single-stranded telomeres, we developed a single-molecule fluorescence resonance energy transfer (smFRET) experiment using a TG12-end substrate. As shown in Fig. 1A, a fluorescence donor (Cy3) and an acceptor (Cy5) are attached to the tailed duplex DNA substrate (TG12-end) separated by a 12-nt single-stranded (ss) DNA overhang of yeast telomeric TG1-3 sequences (Fig. S1). In the TG12-end substrate alone, the flexible ssDNA brings Cy3 and Cy5 dyes in proximity, resulting in a high FRET state (FRET ∼ 0.74, 50 mM NaCl, Fig. 1B). The physiological intracellular ionic concentration in yeast has been estimated to range widely, from less than 20 mM to ∼200 mM K^+^ ions (22–24). Therefore, we also carried out the experiments using 150 mM KCl, resulting in FRET ∼ 0.87 (150 mM KCl, Fig. 1B). FRET efficiency is highly sensitive to the distance between the dye pair and the relative orientation of their dipole moments. When proteins bind to the ssDNA segment between the dye pair, they may alter the local photophysical microenvironment, their relative orientations, or the flexibility of the ssDNA segement, thereby influence FRET efficiency. Thus, FRET experiments can be used to monitor changes in the DNA configuration associated with Cdc13 binding.

**Figure 1.**
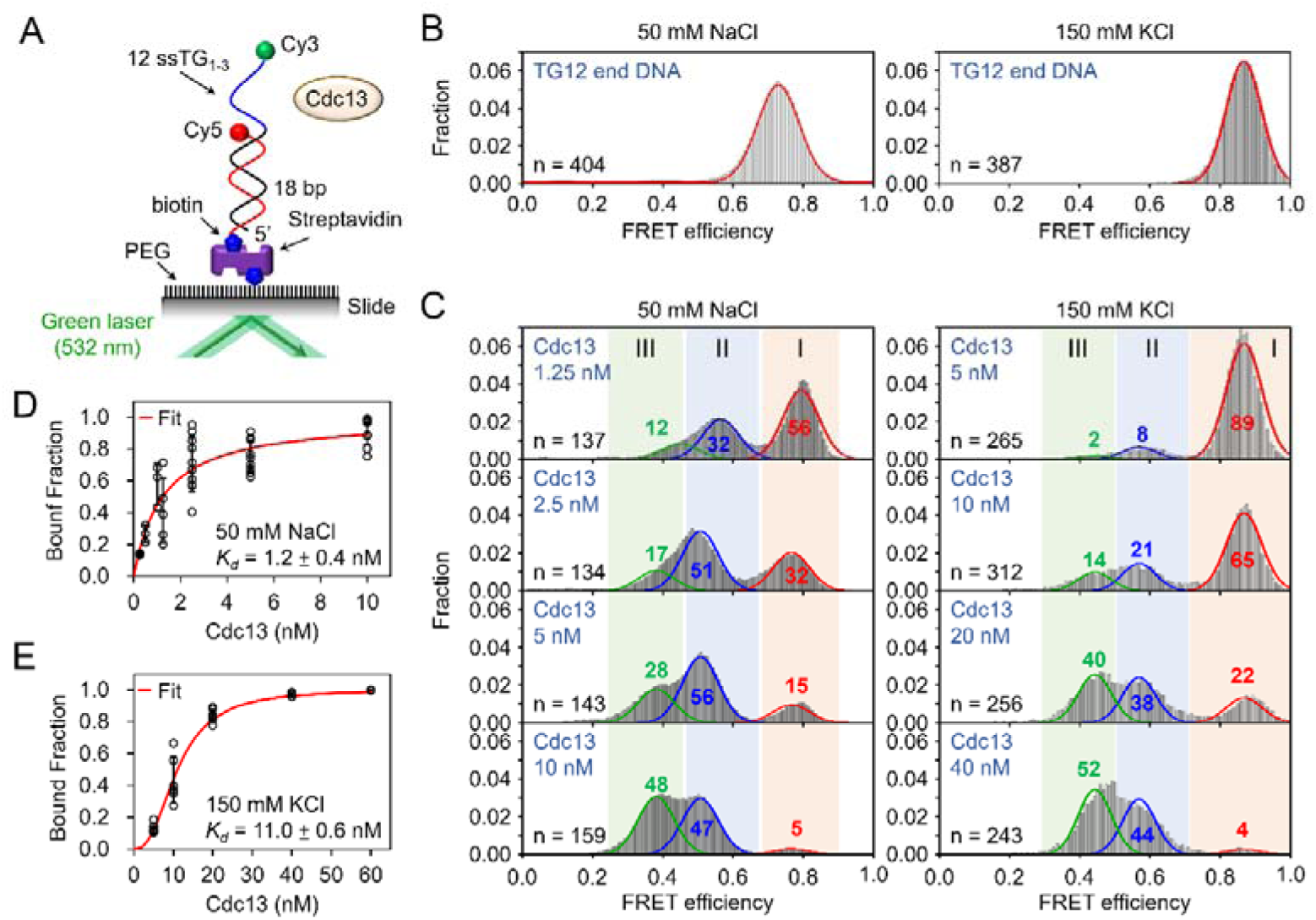
smFRET reveals two Cdc13-DNA binding states. **(A)** smFRET experimental setup. Biotin-PEG-coated microscope slides are loaded with streptavidin and anchored with the tailed-duplex TG12-end DNA. The FRET donor Cy3 and acceptor Cy5 are colored in green and red, respectively. Binding of Cdc13 to this DNA substrate causes decreases in FRET efficiency. **(B)** TG12-end DNA returns with a FRET efficiency of 0.73±0.06 (50 mM NaCl) or 0.85±0.12 (150 mM KCl), with n denoting numbers of analyzed molecules. **(C)** Histograms of FRET efficiency distributions of TG12-end DNA in the presence of various Cdc13 concentrations. Histograms of 50 mM NaCl were fitted by three Gaussians (red curves), with peak values of 0.74±0.07 (state I, red shading), 0.48±0.07 (state II, blue shading) and 0.35±0.04 (state III, green shading). Histograms of 150 mM KCl were fitted by three Gaussians (red curves), with peak values of 0.85±0.12 (state I, red shading), 0.57±0.12 (state II, blue shading) and 0.45±0.12 (state III, green shading). Numbers within the peaks indicate the population percentages. Data were taken 3 mins after Cdc13 addition. **(D)** Binding curve of Cdc13 on TG12-end DNA at 50 mM NaCl. Bound fraction is the sum of state II and III. The apparent dissociation constant (*K_d_*) is presented as value±SEM. Fitted binding curve are presented. Error bars represent standard deviations. **(E)** As in **(D)**, binding curve of Cdc13 on TG12-end DNA at 150 mM KCl.

Recombinant budding yeast Cdc13 isolated from insect cells was prepared to high purity (Fig. S2). Cdc13 binding to TG12-end substrate resulted in the decrease of the DNA-only population (Fig. 1C) and, surprisingly, the appearance of two additional lower FRET states with efficiency centered at ∼0.48 (state II, blue shading) and at ∼0.35 (state III, green shading), for 50 mM NaCl, and ∼ 0.57 and ∼ 0.45 for 150 mM KCl. Each FRET state can be well fitted by a Gaussian distribution, with the area of each Gaussian representing its relative population. The population of each FRET state is indicated in the corresponding FRET histograms, as shown in Fig. 1C. As Cdc13 concentration increased, a quick increase in state II population and a slower delayed increase in state III population can be seen at both salt conditions (Fig. 1C). Two distinct lower FRET states indicate two DNA states upon Cdc13 binding. State III appeared to represent the final reaction product, as increasing Cdc13 concentration up to 160 nM shifted the population from state II to state III (Fig. S3A). Under both salt conditions, Cdc13 exhibits a similar binding pattern.

As the presence of FRET states II and III depends on Cdc13, we defined the Cdc13-bound fraction as the sum of the fractions of FRET states II and III. Analysis of the Cdc13 binding curve to TG12-end telomeric DNA yields an apparent *K_d_* of 1.2 ±0.4 nM (Fig. 1D, 50 mM NaCl) and 11.0±0.4 nM (Fig. 1E, 150 mM KCl). Similar binding was observed in a tailed duplex substrate carrying a 15-nt telomeric ssDNA tail (TG15-end, Fig. S3B). Cdc13 can also associate with Stn1 and Ten1 to form the CST complex (25). We also purified individual Stn1 and Ten1 proteins, but including them did not appear to affect formation of state II and III (Fig. S4).

To investigate whether the formation of FRET states II and III requires specific binding of Cdc13 to telomeric DNA, we used a tailed-duplex DNA substrate with a 13-nt polyT overhang (T13-end DNA). This substrate showed no significant change in FRET efficiency (Fig. S5A), suggesting that the FRET changes observed with telomeric DNA depend on the specific DNA-binding properties of Cdc13 seen in Fig. 1C.

To determine whether Cdc13 binds or associates the duplex DNA segment in the TG12-end substrate, we placed Cy3 fluorophore positioned within the duplex region (TG12 int-5 and TG12 int-8, Fig. S5B-C and Fig. S1). Addition of Cdc13 did not affect the FRET efficiency of either substrate, consistent with previous studies (1, 2). These results indicate that two additional distinct FRET states seen in TG12-end DNA result from Cdc13 binding to telomeric ssDNA.

### FRET state III is less susceptible to dissociation

To test the stability of the Cdc13-bound FRET states II and III, the Cdc13-DNA complexes were challenged with increasing salt concentrations of NaCl. As protein-DNA binding typically involves electrostatic interactions, a stable protein-DNA complex may be more resistant to elevated salt concentrations. We first populated FRET states II and III using high concentrations of Cdc13 in a buffer containing 50 mM NaCl. We then challenged the Cdc13-DNA complex with buffer containing 50, 75, or 100 mM NaCl (Fig. S6 and Figure 2A). Surprisingly, the state III fractions were not changed upon buffer washes at these NaCl concentrations, while the state II fractions were significantly dropped at NaCl concentrations above 50 mM with a concomitant increase in state I. Thus, FRET state III is less susceptible to dissociation, reflecting its higher stability and potential biochemical roles.

**Figure 2.**
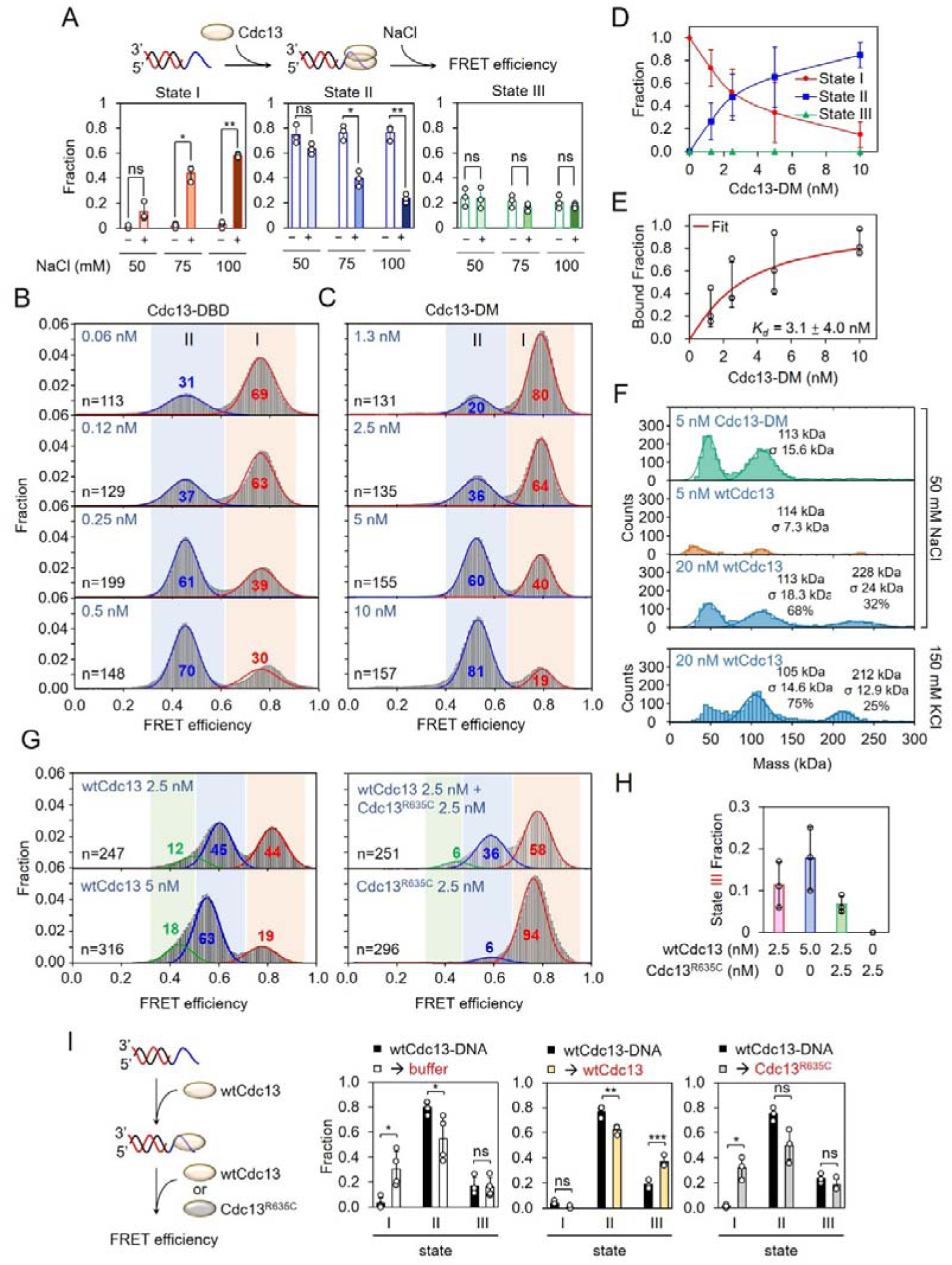
FRET state III is salt resistant and its formation requires Cdc13’s DNA binding and dimerization activity. **(A)** Salt challenging experiments were done by incubating Cdc13 on TG12-end DNA first in buffer containing 50 mM NaCl, followed by challenging with 50, 75, and 100 mM NaCl. Fractions of FRET state I, II and III before (-) and after (+) the salt challenge are shown. Averages of 3 independent experiments are presented. Error bars represent standard deviations. Paired and two-tailed Student’s t-test was applied to assess whether the means of the two groups were statistically different from each other, with * denoting p<0.05 and ** denoting p<0.01. ns, not significant. **(B)** FRET histograms of 0.06, 0.12, 0.25, and 0.5 nM Cdc13-DBD incubated with TG12-end DNA are presented. Cdc13-DBD cannot form state III. **(C)** FRET histograms of 1.3, 2.5, 5, and 10 nM Cdc13-DM incubated with TG12-end DNA are presented. Dimerization-defective Cdc13-DM mutant fails to form state III. **(D)** Fractions of state I (red), II (blue), and III (green) at different Cdc13-DM concentrations. No state III is populated. **(E)** Binding curve of Cdc13-DM mutant on TG12-end DNA. Bound fraction is the sum of state II and III. Fitted binding curve are presented. Error bars represent standard deviations. **(F)** Mass photometry analysis of Cdc13 and Cdc13-DM oligomerization state. Solid curves represent Gaussian fits to the experimental histograms. Measured molecular masses, standard deviations (σ), and relative population percentages for monomeric and dimeric species are indicated above each peak. Low molecular weight peaks below 100 kDa (around 50 kDa) represent background or noise particles. **(G)** FRET histogram of 2.5 nM wtCdc13, 5 nM wtCdc13, 2.5 nM Cdc13^R635C^, mixture of 2.5 nM wtCdc13 and 2.5 nM Cdc13^R635C^ with TG12-end DNA. **(H)** Comparison of state III fraction at various wtCdc13 and Cdc13^R635C^ mutant concentrations. Averages of 3 independent experiments are presented. Error bars represent standard deviations. **(I)** Cdc13 chasing experiments were done by incubating wtCdc13 on TG12-end DNA first, followed by challenging with buffer-only, wtCdc13 or Cdc13^R635C^ mutant. Fractions of FRET state I, II and III before (filled black bars) and after (color-indexed bars) the challenge are shown. Averages of 3 independent experiments are presented. Error bars represent standard deviations. Paired and two-tailed Student’s t-test was applied to assess whether the means of the two groups were statistically different from each other, with * denoting p<0.05, ** denoting p<0.01, and ns denoting not significant.

### Dimerization and DNA-binding of Cdc13 required for state III formation

Cdc13 includes several functional domains, such as DNA-binding and dimerization domains. For example, Cdc13 can form dimers in solution (13,16,26). Four mutations in the OB1 domain have previously been shown to disrupt Cdc13 dimerization (13). To determine which domains are responsible for the observed FRET states II and III, we purified the DNA-binding domain of Cdc13 (Cdc13-DBD), and the dimerization-defective mutant (Cdc13-DM) (Fig. S2). With the increasing Cdc13-DBD concentration, the apparent increase in the FRET state II population was detected, but no FRET state III was seen (Fig. 2B). This result indicated that Cdc13’s DNA-binding domain only, in the absence of the dimerization domain, is not sufficient to populate FRET state III. Similarly, dimerization-defective Cdc13-DM mutant induced the formation of state II populations in a dose-dependent manner (Fig. 2C-D), but it failed to populate FRET state III. Interestingly, the apparent DNA binding affinity of the Cdc13-DM mutant (*K_d_* ∼3.1 nM) was similar to that of the wild type. (Fig. 2E). Thus, dimerization of Cdc13 is required for state III formation, based on the results from DNA-binding domain only (Cdc13-DBD) and dimerization-defective (Cdc13-DM) mutants.

Results from Cdc13-DM studies suggest that state II represents binding of monomeric Cdc13 to DNA. We used mass photometry measurements to verify the presence of Cdc13 monomer (Fig. 2F). Cdc13 is a 924-amino-acid protein with a molecular mass of 104,895 Da (14). Cdc13-DM was confirmed as a single monomeric 113 kDa species in mass photometry. At 5 nM wtCdc13, monomeric Cdc13 is the dominant population in solution. At 20 nM wtCdc13, two populations were observed: one at 113 kDa and the other at 228 kDa, suggesting the formation of dimeric protein in solution containing 50 mM NaCl. A similar distribution was also observed at 150 mM KCl. The results indicate that Cdc13 exists in a concentration-dependent monomer-dimer equilibrium in solution across different ionic conditions, with the monomer predominating at low protein concentration, consistent with monomeric Cdc13 binding to DNA in state II.

Previous studies showed that a single R635C mutation of Cdc13 is defective in DNA binding but sufficient in dimer formation (27). As expected, Cdc13^R635C^ mutants failed to form the FRET state II due to the lack of DNA-binding properties (Fig. S7). Next, the mixture of Cdc13^R635C^ mutant and wtCdc13 was used to validate the observation of monomer-dimer equilibrium in mass photometry (Fig. 2G-H). If there are monomer-dimer exchange between wtCdc13 and Cdc13^R635C^, effective wtCdc13 monomer and dimer would decrease, which led to reduced fraction of both state II and III. As a control, wtCdc13 concentration was increased from 2.5 nM to 5.0 nM, which increased state III fraction from 0.11 to 0.18 as previous described (Fig. 1C). However, in the mixture of 2.5 nM wtCdc13 and 2.5 nM Cdc13^R635C^ mutant, the state III fraction reduces to 0.05, even lower than the 2.5 nM wtCdc13 alone. The experimental observation was consistent with the hypothesis that wtCdc13 and Cdc13^R635C^ form mixed dimers in solution, thereby reducung the availability of wtCdc13 for DNA binding.

To validate whether additional Cdc13 with DNA-binding activity is required for the state III, we designed a chase experiment (Fig. 2I). We first prepared states I/II/III using 10 nM wtCdc13, and then chased them with additional 8 nM wtCdc13 or Cdc13^R635C^ mutant. Because the second Cdc13 molecule accommodated in TG12-end DNA may increase steric hindrance and thereby reduce FRET efficiency, we predicted that adding Cdc13 would increase the fraction of state III, which corresponded to the lower FRET efficiency population. Our single-molecule experimental setup allowed efficient buffer exchange for the surface-anchored protein-DNA substrates, making it possible to monitor the fraction change of these chase experiments. Control experiments using buffer only during the chase (wtCdc13-DNA buffer) returned a slight drop in state II and a corresponding increase in state I, with no change in the fraction of the state III. This is consistent with the 50 mM NaCl wash shown in Fig. 2A. If the state II to state III conversion requires an additional Cdc13 with DNA-binding property, the predictions are: (1) the chase with additional wtCdc13 will shift state II into state III, leading to the increased fraction of state III, and (2) the chase with Cdc13^R635C^ mutants show no change in state III fractions, but resulting in a significant drop in state II. Our experimental observations are consistent with these predictions (Fig. 2I, Fig. S8A). Moreover, real-time experiments observed obvious higher transition events of state II to state III in wtCdc13, while no such were observed seen in Cdc13^R635C^ mutant (Fig. S8B-C). Together, these results supported a requirement for DNA binding activity of the incoming Cdc13 molecule during the conversion of II III.

### Two Cdc13 molecules are found in TG12-end DNA

The formation of FRET state III requires Cdc13’s dimerization and DNA-binding properties. To clearly determine the number of Cdc13 molecules bound to TG12-end DNA, we designed the colocalization single-molecule spectroscopy (CoSMoS) experiments to directly visualize the numbers of dye-labeled Cdc13 molecules bound to DNA. This technique takes advantage of the multi-color fluorescence colocalization of proteins and individual DNA molecules (28), and has been applied to study the mechanism of transcription initiation at a promoter (29), and replication initiation (30). At the single-molecule fluorescence level, each individual fluorescence exhibits a single-step photobleaching. Combining CoSMoS with the single-molecule photobleaching experiments allows us to determine the stoichiometry of proteins within a complex bound to DNA (31,32). A single DY-649P1-labeled peptide was covalently linked to individual Cdc13 molecules using the sortase labeling strategy (Fig. S9A-C) (33,34), with a labeling efficiency of ∼75%. In our experiment, a Alexa488-labeled TG12 DNA was tethered to a microscope slide (Fig. 3A), and the colocalization of Alexa488-DNA and DY-649P1-Cdc13 signals can be readily detected at the single-molecule level (Fig. 3B). We confirmed that DY-649P1-labeled Cdc13 showed an apparent *K_d_* value of 0.4±0.04 nM (Fig. 3B), similar to the affinity determined in FRET-based assays using no-fluorescence-labeled wtCdc13 (Fig. 1D). Controlled experiments using DY-649P1-Cdc13 and T13-end ssDNA substrate returned with no apparent binding, as expected.

**Figure 3.**
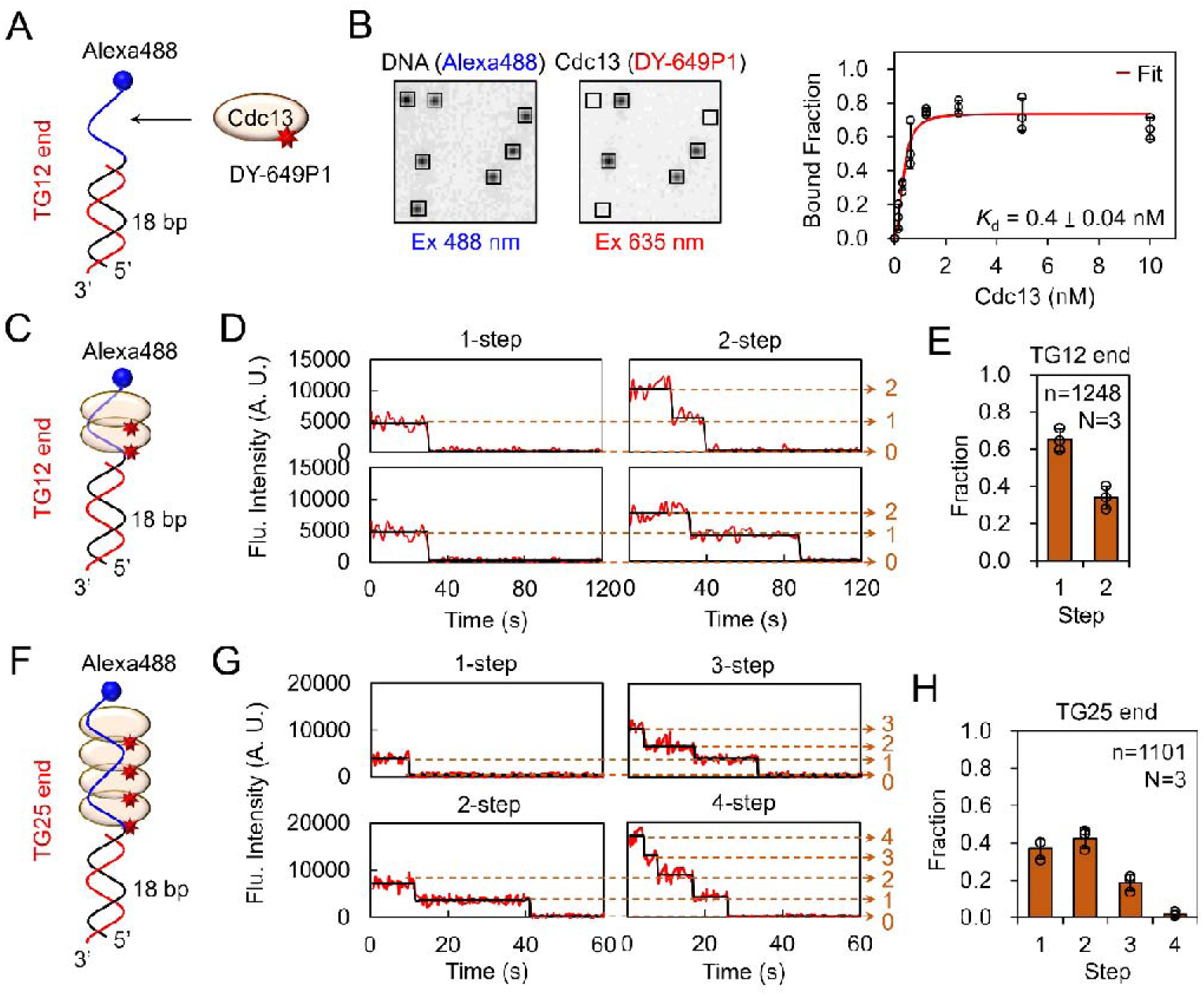
Counting numbers of DY-649P1-labeled Cdc13 monomers at individual telomeric DNA substrates. **(A)** CoSMoS experimental setup. A TG12-end DNA labeled with biotin at one end and Alexa488 at the 3’-terminating telomeric end. DY-649P1 labeled Cdc13 is then added to the DNA and fluorescent signals are observed by TIRF microscopy. **(B)** Exemplary images when excited with 488 nm and 635 nm lasers to visualize the DNA (Alexa488, left) and Cdc13 (DY-649P1, right). Squared boxes were individual fluorescence spots scored in the DNA channel and were mapped into the DY-649P1 channel. 4 out 7 DNA molecules are colocalized with DY-649P1-labeled Cdc13. Fractions of colocalized DY-649P1-Cdc13-DNA spots at different Cdc13 concentrations on TG12-end DNA (red) are shown. Error bars represent standard deviations. **(C)** Photobleaching steps analysis of TG12-end DNA-bound DY-649P1-Cdc13. 10 nM of DY-649P1-Cdc13 was first incubated with TG12-end DNA substrates, and then washed with buffer to remove unbound Cdc13 to proceed photobleaching. **(D)** Examples of photobleaching steps in TG12-end DNA showed both the one- and two-step photobleaching events. Each photobleaching step represents the presence of a single DY-649P1-labeled Cdc13. Note the intensity of one DY-649P1-Cdc13 is around 5000. **(E)** Fraction of 1-step and 2-step photobleaching events in TG12-end DNA. Results were obtained from 3 independent experiments, monitoring a total of 1248 molecules. At most, two-step photobleaching is seen in TG12-end DNA. **(F-H)** As in **(C)-(E)**, but using DNA carrying 25-nt single-strand telomeric DNA (TG25-end). Examples of 1-4 photobleaching steps are seen **(G)**. Fractions of photobleaching steps **(H)** with data from 3 independent experiments, monitoring a total of 1101 molecules. At most, 4-step photobleaching can be seen in TG25-end DNA.

After incubation of 10 nM of DY-649P1-Cdc13 on surface-bound DNA, buffer washes were applied to remove free, unbound DY-649P1-Cdc13. Photobleaching experiments were then conducted to determine the stoichiometry of DY-649P1-Cdc13 binding to TG12-end DNA (Fig. 3C). Only DY-649P1-Cdc13 signals colocalized to the Alexa488-labeled TG12-end DNA substrates were analyzed. Clear 1-step or stepwise 2-step photobleaching of DY-649P1-Cdc13 can be seen (Fig. 3C-D). Note that intensities of DY-649P1 before photobleaching are similar and scalable, with ∼4500 for 1-step and ∼9500 for 2-step. Among a total of 1248 colocalized DY-649P1-Cdc13/Alexa488-DNA molecules analyzed out of 3 independent experiments, 65% of the colocalized Cdc13-DNA molecules showed one-step DY-649P1 photobleaching and ∼34% showed two-step photobleaching (Fig. 3E). Photobleaching experiments directly indicated that at most two Cdc13 molecules are bound to the TG12-end DNA. We also compared the experimental photobleaching-step distribution with a binomial model acoounting for the labeling efficiency, which assuming 1-5 Cdc13 molucules per DNA molecule. TG12 experimental distribution showed the best agreement with the simulation assuming two Cdc13 molecules per DNA molecule (Fig.S9D). After acoounting for the ∼75% DY649P1 labeling efficiency. The estimated fraction of DNA-bound Cdc13 dimer was ∼54%. This extimate was also comparable to the fraction observed in FRET experiments (Fig. 1C).

We also used a longer telomeric ssDNA tail carrying 25-nt in the analysis. Using this TG25-end DNA substrate (Fig. 3F), we observed photobleaching steps as many as 4, with exemplary photobleaching traces presented in Fig. 3G. Among a total of 1101 colocalized DY-649P1-Cdc13/Alexa488-DNA molecules analyzed out of 3 independent experiments, fractions of ∼0.37, ∼0.42, ∼0.19, and ∼0.02 of molecules were photobleached in 1-, 2-, 3-, and 4-steps, respectively for TG25-end DNA (Fig. 3H). Consistently, TG25 experimental distribution showed the best agreement with the simulation assuming four Cdc13 molecules per DNA molecule (Fig.S9D).

### Dynamic interchange among different Cdc13-DNA bound states

The FRET and CoSMoS experiments allow the monitoring of DNA conformational change induced by Cdc13 binding to telomeric ssDNA in real-time. For example, upon the Cdc13 addition to TG12-end substrate (the shaded area around 50 s), the dynamic transitions among three FRET states were observed (Fig. 4A). The exemplary time course showed a sequential conversion from state I (DNA-only) to state II right after the deadtime and then transition to state III. We also used the colocalized CoSMoS signal to directly show the binding of individual DY-649P1-Cdc13 on DNA (Fig. 4B). DY-649P1-Cdc13 was added at time zero, and sequential binding of the first and the second Cdc13 molecules can be seen in the exemplary time course (Fig. 4B). For real-time FRET measurements, at each Cdc13 concentration, about ∼200-400 of these transition events were observed from ∼200-300 molecules. More than 50% of these observed transition events were state I II transitions (solid circles, Fig. 4C), which is consistent with the oligomeric observation in mass photometry (Fig. 2G). Notably, with increasing Cdc13 concentration, apparent transition events of state II III and I III were increased. Only a few transition events were observed to initiate from state III (III I and III II), confirming that state III represents a stable Cdc13-DNA complex. The percentages of transition events corresponding to apparent Cdc13 association (top, Fig. 4C, I II, II III and I III) and dissociation (middle, Fig. 4C, II I, III II and III I) are shown. Data from FRET experiments are displayed in the unshaded regions and those from CoSMoS are in the gray-shaded regions. To minimize background fluorescence in real-time detection of individual protein-binding events, CoSMoS measurements were conducted at lower Cdc13 concentrations. We also used FRET to monitor Cdc13 binding to TG12-end telomeric ssDNA at 150 mM KCl (Fig. 4C, bottom; Fig. S10). Similarly, transitions from state I to II predominated at 150 mM KCl. The exemplary time trace also showed sequential transitions from state I to state II and subsequently to state III at 150 mM KCl (Fig. S10). Consistent with the higher apparent *K_d_* of wtCdc13 at 150 mM KCl (Fig. 1E), frequent transitions between state I and state II were observed.

**Figure 4.**
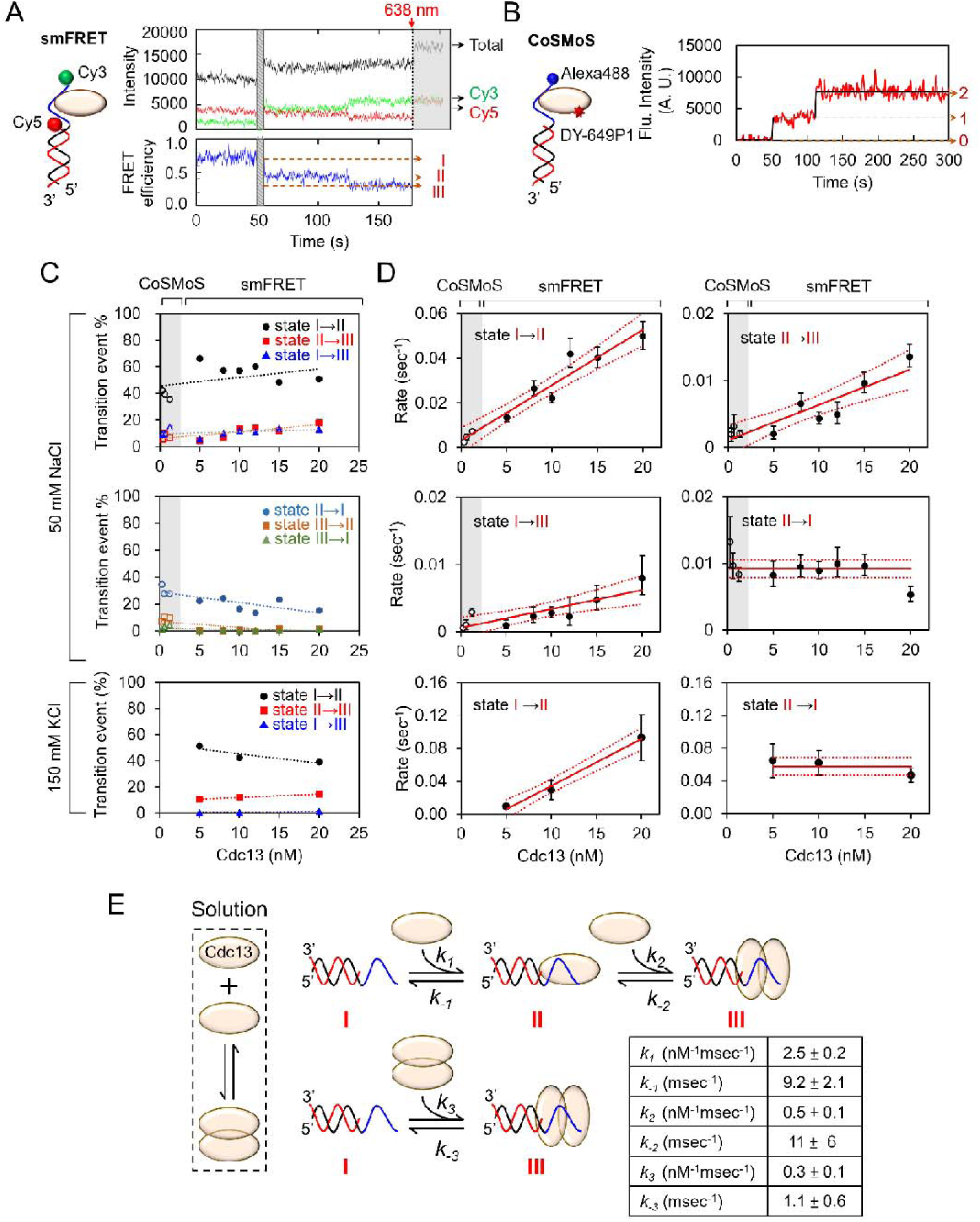
Kinetic analysis of FRET and CoSMoS experiments identify sequential Cdc13 monomer binding. **(A)** Representative single-molecule FRET time-course showing dynamic conversion among FRET states I, II, and III upon Cdc13 binding to TG12-end DNA. Total fluorescence, donor Cy3, and acceptor Cy5 intensities are colored in black, green, and red, respectively. Corresponding FRET efficiencies are shown below in blue. Dashed red lines correspond to three FRET states. The 638 nm laser was turned on at t = 175 sec (red arrow) to ensure the presence of Cy5 at the end of each experiment. Schematics on the left show the position of the fluorophores in single-molecule FRET. **(B)** Representative CoSMoS time-course showing dynamic binding of DY-649P1-labeled Cdc13 colocalized with Alexa488-TG12-end DNA. Binding and dissociation of DY-649P1-Cdc13 can be identified in steps, with at most two Cdc13 molecules bound. In this real-time binding experiments, 0.3, 0.6 and 1.25 nM of DY-649P1-labeled Cdc13 concentrations were used. Schematics on the left show the position of the fluorophores in CoSMoS. **(C)** Percentages of transition events from both CoSMoS (open symbols) and FRET (filled symbols) time-courses. Association events at 50 mM NaCl (top), dissociation events at50 mM NaCl (middle), and association events at 150 mM KCl (bottom). Even with the different experimental designs of CoSMoS and FRET, as well as different Cdc13 concentrations used, both sets of data align for the same trend. For each Cdc13 concentration, over 200 (smFRET 200-400; CoSMoS 200-900) transition events were determined (N≧3). **(D)** Transition rates from both CoSMoS (open symbols) and FRET (filled symbols) time-courses between Cdc13-TG12-end DNA binding events. State time-coursed determined by vbFRET were analyzed by the homogeneous Markov model to return transition rates at each Cdc13 concentrations. Note that data from CoSMoS and FRET align well for each transition. For association transitions, rates are linear correlated with Cdc13 concentrations. The 95% confidence intervals for the rate estimates are shown as red dashed lines around the fitted trend line. Error bars represent the sample standard deviation. **(E)** Cdc13 sequential binding model and the rate constants determined from 50 mM NaCl are presented.

Dynamic switching among these FRET states can be scored and fitted by the hidden Markov modeling (HMM) method (35). Dwell times associated with each state are determined from the HMM analysis. Time series of these assigned states and their transition events at given Cdc13 concentrations allow the determination of apparent rate constants for transitions among these three FRET states. We then assumed the reaction kinetics by a 3-state continuous homogeneous Markov model to estimate the transition rates that maximize the likelihood of all state time series (see Methods for details).

The rates that best describe all FRET and CoSMoS time series based on the kinetic model are represented in the prevalence plots (Fig. S11 and Fig. S12 for 50 mM NaCl and 150 mM KCl, respectively) and with rates presented in Fig. 4D (solid circles for FRET and open circles for CoSMoS) for each Cdc13 concentration analyzed. Note that large uncertainties are seen in *k_-2_* and *k_-3_* due to the very limited observed dissociation events from state III (III II and III I, Fig. 4C middle, Fig. S13). We presented the all rate constants in Fig. 4E (50 mM NaCl). Apparently, linear Cdc13 concentration-dependence can be seen for association reactions (I II, II III and I III), and the dissociation reaction of II I does not depend on Cdc13 concentration, as expected (Fig. 4D). Surprisingly, data from CoSMoS experiments (open symbols) correlate excellently with the FRET experiments (filled symbols), considering different experimental layout and Cdc13 concentration range used. As the same set of kinetics parameters can be used to describe the conversion of FRET states and DY-649P1-Cdc13 bindings, FRET state II is biochemically the same as one Cdc13 binding and FRET state III is two Cdc13 binding. In our kinetic model, both sequential Cdc13 monomer binding (I II III) and direct Cdc13 dimer binding (I III) are possible, as both are seen experimentally. As shown in Fig. 4E, the rate constant (slopes) of sequential binding is 8-fold higher than the direct dimer binding (*k_1_* =2.5, *k_3_* =0.3 nM^-1^ msec^-1^), making sequential binding kinetically favored.

In addition, time-courses of the Cdc13-DM mutant only showed transitions between FRET state I and II (Fig. S14). Kinetic analysis of these Cdc13-DM mediated transitions returns with rate constants of *k_1_* (I II) and *k_-1_* (II I), as shown in Fig.S14. These rate constants are, in general, the same as the wtCdc13 (Fig. 4E): *k_1_* of 3.7±1.3 nM^-1^ msec^-1^, and *k_-1_* of 8.9±5.9 msec^-1^ for Cdc13-DM (Fig. S15). These similar kinetics parameters involved in the conversion between FRET state I and II suggested that the DNA binding domain of Cdc13 is responsible for populating FRET state II. As Cdc13-DM mutants failed to populate FRET state III, it reiterates the need for a Cdc13 dimerization domain for state III.

## Discussion

The yeast *Saccharomyces cerevisiae* carries a 12-14 base single-stranded telomeric tail at the ends of its chromosomes throughout most of the cell cycle, except during the S phase when telomeres are replicated (20). In cells, this 12-14 nt long telomeric tail is bound by telomere-specific binding proteins Cdc13 to protect the telomere and to coordinate telomere length maintenance. Here, we use two complementary single-molecule experiments (FRET and CoSMoS) and an *in vitro* reconstituted system to characterize how Cdc13 binds to telomeric DNA. We show that Cdc13 monomer kinetically favors binding to telomeric DNA sequentially to form Cdc13 dimers on DNA with higher stability. Cdc13 dimers can also bind directly to telomere DNA in parallel. The formation of the Cdc13 dimer-DNA complex requires both dimerization and DNA-binding activities of the protein, although the temporal sequence of dimerization and DNA-binding could not be distinguished now. The CoSMoS analysis of colocalized dye-labeled Cdc13 and DNA allows direct validation of two Cdc13 molecules binding to telomeres. With the combined results from single-molecule FRET providing information on DNA conformation and CoSMoS dictating the numbers of Cdc13 binding, we provide the key mechanistic details responsible for the stable Cdc13’s binding events.

Substantial evidence supported the formation of Cdc13 dimers in solution (13,16,17) making it widely assumed that Cdc13 forms dimers in solution and binds directly to telomeres. Our results support a novel Cdc13-binding mechanism, in which monomeric Cdc13 binds sequentially to telomeric DNA and forms a stable dimer-DNA complex at low local Cdc13 concentration (e.g. in G1 phase). At higher local Cdc13 concentration (e.g. in S phase), the complex may also form through the binding of preassembled solution dimers or the rapid recruitment of a second monomer. The Cdc13 dimer-DNA complex is resistant to washing at elevated salt concentrations, suggesting its potential regulatory roles in preventing telomeres from nucleolytic degradation and in interacting with other telomere-associated proteins (such as Stn1/Ten1, Est1, and Pol α). Molecular genetic analysis has shown that yeast cells carrying the cdc13-DM mutant fails to interact with DNA polymerase α, do not protect telomeres at high temperature, and possess short telomeres (13). These results suggest that while Cdc13 monomer-DNA binding is kinetically favored, it is not sufficient to protect telomeres and maintain telomere functions. A comparable example is the yeast MCM complex during replication initiation. When MCM is loaded onto replication origins together with ORC and Cdc6, the resulting OCM complex becomes stable and resists to high-salt challenges, allowing this complex to remain stably bound to DNA until replication initiation. This stability is critical for ensuring precise control of replication timing and preventing re-replication during the cell cycle (36,37).

Compared with a direct dimer binding, a sequential monomeric Cdc13 binding to telomeres can offer several regulatory advantages. Compared with the sequence-non-specific single-stranded DNA binding protein RPA, Cdc13 specifically binds to telomeres to execute its function in telomere maintenance. Initial monomeric Cdc13 allows efficient off-target dissociation to achieve increased specificity at the telomere. The binding of the first Cdc13 monomer may allow the creation of a foothold on DNA. A “kinetic proofreading-like mechanism” may be possible, given that the first monomer can quickly dissociate, evident by a large *k_-1_* estimated here. This mechanism allows complex formaiton to be regulated in response to cellular cues to prevent unnecessary or premature complex formation. Sequential binding often requires lower initial binding energy cost and concentration thresholds and offers a more flexible, sequence-specific regulation. Given the kinetically derived *K_d_* for the first monomeric Cdc13 binding is 3.7 nM (the ratio of *k_-1_* and *k_1_*), compared to the apparent *K_d_* is 1.2±0.4 nM (Fig. 1D), this sequential binding could be particularly important for telomere maintenance, where precise recognition of telomeric sequences is crucial for preventing chromosome end degradation and fusion. Sequential binding of protein monomers to DNA has been identified is a useful regulatory strategy. For example, the prokaryotic LexA binds to DNA as a monomer first, and dimerization with a second LexA occurs on the DNA binding site to coordinate SOS response (38). Eukaryotic transcription factors Fos and Jun form a heterodimeric complex on DNA to mediate transcription activation. Interestingly, although these two proteins rapidly form a heterodimer in solution, Jun binds DNA first and then assembles into the final complex with Fos (39). Importantly, this sequential monomer-binding allows for the enhancement of specificity prior to reaching equilibrium (40). Other examples include blue light sensor protein EL222 (41) and ETS family transcription factor PU.1 (42). All proteins bind to DNA as monomers first to scan for specific target sites and later form dimers or oligomers for function.

This study highlighted that DNA-binding activities of both Cdc13 molecules in dimers are required for the stable complex. Since one Cdc13 molecule is sufficient to interact with ∼11 bases of telomeric DNA (19), the mechanism underlying the requirement for both binding activities of Cdc13 molecules remains unclear. The observation of a decreased FRET efficiency (state III) when the second Cdc13 binds to DNA suggests that binding of the second Cdc13 molecule may induce a conformational change to the Cdc13 monomer-DNA complex, allowing two Cdc13 molecules to bind to the DNA.

Given that Cdc13 binds to the 11-base telomeric DNA with differential affinities, where bases located at the 3’ end are not as critical as those at the 5’ end for binding affinity and specificity (43), this conformation change may involve partial rearrangement of the first Cdc13 on DNA to allow subsequent binding of the second Cdc13 on DNA. We further used Alphafold3 to predict a complex containing two Cdc13 molecules and TG12-end DNA (Figure S2C). The model suggests both DBDs can simultaneously engage the 12-nt ssDNA segemnt through distinct yet partially overlapping DNA-binding interfaces, while their dimerization domains interact with each other. The flexible linker connecting the DBD and dimerization domain may allow relative movement between these two domains. This model provides a possible explanation for how two Cdc13 molecules can stably associate with TG12-end DNA.

Importantly, protein binding-induced conformational changes have been observed in other single-strand binding proteins, as seen in Replication Protein A (RPA). RPA is a heterotrimeric protein composed of three subunits, RPA70, RPA32, and RPA14 (44), and RPA contains multiple OB folds to bind ssDNA with different binding modes of different bound lengths. Each OB fold can dynamically bind and dissociate to DNA, but overall RPA stays bound to DNA to regulate the accessibility of single-stranded DNA by other proteins. Interestingly, these different binding modes appear to proceed sequentially, progressing from a low-affinity binding mode to a stable, high-affinity mode (45,46). Further structural and imaging research to determine how the Cdc13 dimer interacts with 12-base telomeric DNA would provide valuable insights into this process.

## Materials and Methods

### Purification of yeast Cdc13 protein

The 6-His tagged Cdc13 was constructed, expressed and isolated from Sf21 cells using baculoviral expression system, as described previously (3).

### DNA substrates

DNA oligonucleotides were purchased from Integrated DNA Technologies (IDT). To anneal smFRET-based tailed duplex DNA, 1 μM of the Cy5-labeled biotinylated oligonucleotides and an excess of Cy3-labeled oligonucleotides were incubated at 85°C for 5 minutes, then cooled to 25°C at a rate of 1°C per minute. For CoSMoS-based tailed duplex DNA, 1 μM of the biotinylated oligonucleotides and an excess of Alexa488-labeled oligonucleotides were annealed using the same program. The annealed DNA was diluted to 10-20 pM using BSA-T50 buffer (0.4 mg/mL BSA or 2 mg/mL BSA, 20 mM Tris-HCl pH 7.4, 50 mM NaCl).

### Single-molecule FRET experiments

An objective-type total internal reflection fluorescence (TIRF) system was used in the smFRET experiments. Lasers of 532 nm (Ventus, ∼30 mW) and 638 nm (Omicron, ∼36 mW) were used for excitation, and a custom-built dual-view system was used in fluorescence imaging. LabVIEW software was used to control the electron-multiplying charge-coupled device (EM-CCD, iXon, Andor).

To tether DNA substrate, the biotin-PEGylated slides were incubated with 0.02 mg/ml streptavidin (SA) at room temperature for 2 minutes, washed three times with BSA-T50 buffer. DNA substrate at 20 pM was introduced, followed by washing with the Image buffer (3.5 mM Trolox, 2 mg/mL BSA, 4 mg/mL glucose, 30 U/mL glucose oxidase, 30 U/mL catalase).

To determine the FRET efficiency of the DNA molecules in the snapshot experiments, 1000 frames of images were taken (50 ms/frame, total 50 sec) with the 532 nm laser excitation. The 638 nm laser was then turned on, and 100 frames were recorded at 50 ms/frame to confirm the presence of Cy5 signal. In real-time FRET experiments, 1000 frames were recorded first for DNA-only, and then Cdc13 was introduced, and 3500 frames were recorded. The 638 nm laser was then turned on at the end of each experiment.

### CoSMoS experiments

A micro-mirror total internal reflection fluorescence (TIRF) microscope (RM21, Mad City Labs) was adapted for CoSMoS experiments and equipped with a piezo stage for precise focus adjustments. Three excitation lasers, 488 nm (Omicron, ∼0.8 mW), 532 nm (MatchBox, ∼1.2 mW), and 635 nm (Coherent, ∼1.5 mW), are used. Fluorescent images were captured with a 100x objective lens (UAPON 100XOTIRF, Olympus) and spatially split using a dual-wavelength separation system (Optosplit II Bypass, Cairn Research) with an FF640-FDi01 dichroic filter (Semrock).

To measure the colocalization (%) of Cdc13 and DNA in snapshot experiments, DY-649P1-Cdc13 was diluted by CoSMoS Image buffer (50 mM Tris-HCl pH7.8, 1 mM EDTA, 50 mM NaCl, 2 mM DTT, 1 mg/mL BSA, 2 mM Trolox, 8 mg/mL glucose, 30 U/mL glucose oxidase, 30 U/mL catalase) to final concentrations of 0.15, 0.3, 0.6, 1.25, 2.5, 5, or 10 nM, applied to DNA substrates at room temperature for 10 minutes. For each field of view, 5-frame images were captured (0.5 sec/frame) to record the DNA locations using a 488 nm laser or 532 nm laser, then switched to a 635 nm laser for another 5 frames (0.5 sec/frame) to record Cdc13 signals. For real-time DY-649P1-Cdc13 binding experiments, DY-649P1-Cdc13 at 0.3, 0.6 or 1.25 nM were applied and recorded a 5-minute movie. Images were captured at 0.2-second exposure time with 488 nm and 635 nm laser in an alternative excitation cycle. Each excitation cycle was composed of 1 frame for each laser excitation.

To determine the stoichiometry of the Cdc13-DNA complex, tethered DNA was incubated with 10 nM DY-649P1-Cdc13 for 10 minutes and then washed with Cdc13 Image buffer. A 2-minute movie was recorded using 488 nm and 635 nm laser. Images were taken at 0.2-second exposure time in an alternating excitation cycle. Each excitation cycle was composed of 1 frame for each laser excitation.

### Mass photometry experiment

Molecular weight and oligomeric state of Cdc13 in solution were collected using a TwoMP Mass Photometer (Refeyn). Bovine serum albumin and thyroglobulin in phosphate buffered saline were used to generate a calibration curve for molecular weight determination. The experiments were perforemed in T50Na (50 mM Tris-HCl pH7.8, 50 mM NaCl) or T150K (50 mM Tris-HCl pH7.8, 150 mM KCl) at room temperature. Samples were analyzed following the manufacturer’s protocols, and the data were graphed using DiscoverMP software or Origin 9 software.

## Acknowledgements

We thank Audra Amasino and Stephen Bell for sharing and discussing the sortase labeling protocols, and Peter Chi for providing cloning fragments. We thank Cheng-Hsin Huang, Jun-Cheng Zhuang, and Richard P. Cheng (NTU Chemistry) for providing HPLC and lyophilization instruments, advice on the optimization of peptide purification protocols, and the help on MALDI analyses. We thank NTU Mass Spectrometry Consortia for providing mass spectrometry instruments and services.

## Funding

This work was financially supported by the “Center of Precision Medicine” from The Featured Areas Research Center Program within the framework of the Higher Education Sprout Project by the Ministry of Education (MOE), and the National Science and Technology Council (111-2311-B-002-009-MY3 and 112-2311-B-002-013 to JJL, and 107-2113-M-002-010-MY3 and 113-2123-M-002-011 to HWL) in Taiwan.

## Conflict of interest

The authors declared that there was no conflict of interest.

## Data Availability

All data are incorporated into the article and its online supplementary material.

## Supplementary Materials and Methods

### Preparation of multilayer PEG slides

Microscope slides were cleaned by sonication in 2 M KOH solution for 20 minutes, followed by sonicating the slides twice with alternating water and 95% ethanol for another 20 minutes. The cleaned slides were rinsed with water and dried with nitrogen gas. These slides were then silanized with 2% 3-aminopropyltriethoxysilane (APTES) (Sigma-Aldrich, cat #A3648-100ML) in the dark for 10 min. PEGylation of the slides was achieved by incubating 240 mg/ml methoxypolyethylene glycol succinimidyl carbonate (mPEG-SC) (Laysan Bio, cat #105-183) and 0.6 mg/ml biotin polyethylene glycol (biotin-PEG) (Laysan Bio, cat #109-62) at room temperature for 4 hours in the dark. The biotin-PEGylated slides were rinsed with water and dried with nitrogen gas.

### Preparation of the GGG-Cdc13 protein

The Cdc13 with three glycines (GGG) at the N-terminus, GGG-Cdc13, was expressed and purified from insect cells using the Bac-to-Bac baculovirus expression system. The plasmid pFastBac1-6-His-SUMO-GGG-CDC13 was constructed through GenBuilder DNA assembly. Briefly, the GGG-CDC13 DNA fragment was PCRamplified from yeast YPH499 genomic DNA, ligated to the 6-His-SUMO DNA fragment from pNRB150-SUMO-scDmc1 (kindly provided by Peter Chi, National Taiwan University, Taipei, Taiwan), and cloned into plasmid pFastBac1. The resulting plasmid enabled the expression of Cdc13 with a 6-His-SUMO fusion at the N-terminus. For baculovirus production, the plasmid was transformed into E. coli DH10Bac strain to generate recombinant bacmid. The bacmid was subsequently transfected into Sf21 cells using Cellfectin II (ThermoFisher) for baculovirus production. The baculovirus was amplified in Sf21 cells and later used to infect High Five cells for the expression of 6-His-SUMO-GGG-Cdc13. Harvested cells were washed with phosphate-buffered saline (PBS) buffer and then stored at -80.

To purify 6-His-SUMO-GGG-Cdc13, approximately 2 x 10^7^ High Five cells were resuspended in NP40 lysis buffer (50 mM Tris-HCl pH 8.0, 150 mM NaCl, 1% Nonidet P-40), supplemented with protease inhibitors (1 mM PMSF, 1x protease inhibitor cocktail, TargetMol). After a 30-min incubation, the cells were lysed by sonication and then centrifuged to collect soluble proteins from the cell extract. Ni-NTA-agarose (Agarose Bead Technologies) and Bio-Rex ion exchange resin (Bio-Rad) were used for the purification of 6-His-tagged SUMO-GGG-Cdc13. The resulting protein was dialyzed into storage buffer (50 mM Tris-HCl pH 7.8, 150 mM NaCl, 1 mM dithiothreitol, 20% glycerol), aliquoted, and stored at -80oC after snap-freezing using a dry ice-ethanol bath.

### Preparation of fluorescently labeled peptide

Cdc13 was fluorescently labeled using sortase-mediated transpeptidation (1), with a protocol modified from a previous publication (2). The H2NCHHHHHHHHHHLPETGG-COOH peptide (>85% purity, Genscript) at ∼3 mM was incubated with 1 mg DY-649P1-maleimide (Dyomics) at room temperature for 2 hours in the dark. The reaction was terminated by freezing and lyophilizing. The lyophilized product was resuspended in water and purified using a Vydac 218TP column (218TP1022, Grace). The isolated labeled peptide was verified by analytical HPLC and MALDI-TOF mass spectrometry. The DY-649P1-labeled peptide was lyophilized and stored in -80°C.

### Preparation of DY-649P1-labelled Cdc13

The reaction for 1 mg Cdc13 was conducted at 37°C for 15 min by incubating 2 μM GGG-Cdc13, 2 μM GST-eSortase A, and 10 μM DY-649P1-labeled peptide in sortase reaction buffer (50 mM Tris pH 7.8, 150 mM NaCl, 10 mM CaCl2). The reaction was terminated by adding EDTA to a final concentration of 45 mM. Subsequently, glutathione-agarose (Agarose Bead Technologies) was employed to remove GST-eSortase. Resins pre-equilibrated with GST binding buffer (10 mM Tris pH 7.8, 150 mM NaCl, 1 mM EDTA, 1 mM β-mercaptoethanol) were added to the reaction and mixed for 1 hour to facilitate binding. The unbound fraction was collected by centrifugation.

To eliminate free DY-649P1-labeled peptide, PD-10 columns pre-packed with Sephadex G200 resins (Pharmacia) were utilized. The unbound fraction was applied to PD-10 columns pre-equilibrated with sortase reaction buffer (without CaCl2), and the flow-through fraction was collected. To separate His-tagged Dy649P1-Cdc13 from unlabeled Cdc13, Ni-NTA-agarose (Agarose Bead Technologies) was employed. The flow-through fraction was mixed with resins pre-equilibrated with Ni-NTA binding buffer (50 mM Tris-HCl pH 7.8, 300 mM NaCl, 20 mM imidazole, 20% glycerol, 5 mM β-mercaptoethanol) and mixed for 1 hour to allow binding. The resins were then washed with Ni-NTA binding buffer containing 40 mM imidazole. Cdc13 was eluted with Ni-NTA binding buffer containing 500 mM imidazole and 1% Tween 20. The fractions containing Cdc13 were combined, dialyzed into Cdc13 storage buffer, aliquoted, and stored at -80 after snap-freezing. Purified DY-649P1-Cdc13 was quantified by SDS-PAGE using serially diluted BSA standard solutions.

To determine the labeling efficiency of the sortase reaction, a second labeling method involving the conjugation of Cy3-maleimide to surface-exposed cysteine residues were applied to Cdc13-DY649P1, as previously described (2). The observed colocalized fluorescent signal fraction (Y_1_) was 0.85, and the fraction of molecules exhibiting two-step photobleaching (Y_2_), as showned in Figure 3E, was 0.34. Let X_1_ (0<X_1_<1) denote the DY649P1 labeling efficiency, and X_2_ (0<X_2_<1) denote the fraction of DNA-bound Cdc13 dimers.

Y_1_=X_1_(1-X_2_) + 2X_1_X_2_(1-X_1_) + X_1_X_1_X_2_ = 0.85

Y_2_ = X_1_X_1_X_2_/Y_1_ = 0.34

Solving this system of equations yields X_1_ ∼0.75 and X_2_ ∼0.54. These results indicate that approximately 54% of DNA-bound Cdc13 molecules were dimers after correcting for the DY649P1 labeling efficiency of approximately 75%.

### Purification of 6-His-Ten1

To purify 6-His-tagged Ten1, a 500 mL culture of E. coli harboring pET6H-TEN1 was grown at 37 °C to an OD600 of ∼0.4, then induced with 0.3 mM IPTG and incubated at 16 °C for 16-18 h. Cells were harvested by centrifugation and lysed by sonication in sonication buffer (50 mM Tris-HCl pH 7.8, 150 mM NaCl, and 10% glycerol), supplemented with protease inhibitors. The lysates were cleared by centrifugation and incubated with Ni-NTA agarose resin (GE Healthcare). The resin was washed with sonication buffer containing 300 mM NaCl and 20 mM imidazole, and the protein was eluted with elution buffer (50 mM Tris-HCl pH 7.8, 5 mM β-mercaptoethanol, 250 mM imidazole, 10% glycerol). Eluted protein was dialyzed against dialysis buffer (50 mM Tris-HCl pH 8.0, 50 mM NaCl, 1 mM dithiothreitol, and 20% glycerol). The dialyzed sample was applied to DEAE Sepharose in dialysis buffer, and unbound fractions containing Ten1 were collected. The final protein was aliquoted and snap-frozen in liquid nitrogen for storage at −80 °C.

### Purification of TAP tagged Stn1

Yeast cells harboring the pRS424-STN1-TAP plasmid were grown at 30 °C in synthetic complete medium without tryptophan and supplemented with 2% raffinose until the OD reached ∼1.0. Expression of TAP-tagged Stn1 was induced with 3% galactose at 25 °C for 24 h. Cells from a 4 L culture were harvested, resuspended in buffer containing 25 mM HEPES pH 8.0, 200 mM KCl, 2 mM MgCl, 0.1 mM EDTA, 0.5 mM EGTA, 1 mM dithiothreitol, and supplemented with protease inhibitors, then lysed using a bead beater. The lysate was clarified by centrifugation and incubated with IgG Sepharose 6 Fast Flow (GE Healthcare) in IgG binding buffer (10 mM Tris-HCl pH 8.0, 150 mM NaCl, 0.1% NP-40, and 1 mM dithiothreitol). The resin was washed with the same buffer, and bound proteins were cleaved on-resin overnight with TEV protease in TEV cleavage buffer (50 mM Tris-HCl pH 7.5, 0.5 mM EDTA, 1 mM dithiothreitol). The eluate was incubated with calmodulin affinity resin (Agilent Technologies) in calmodulin binding buffer (10 mM Tris-HCl pH 8.0, 150 mM NaCl, 1 mM Mg-acetate, 1 mM imidazole, 2 mM CaCl, 0.1% NP-40, 10 mM β-mercaptoethanol), followed by washing with binding buffer supplemented with 0.2% NP-40. Bound proteins were eluted using calmodulin elution buffer (10 mM Tris-HCl pH 8.0, 150 mM NaCl, 1 mM Mg-acetate, 1 mM imidazole, 2 mM EGTA, 0.2% NP40, 10% glycerol, 10 mM β-mercaptoethanol), applying a gradient from 2 mM to 10 mM EGTA. Pooled fractions were dialyzed against buffer containing 10 mM Tris-HCl pH 8.0, 100 mM NaCl, 1 mM EDTA, 1 mM dithiothreitol, 40% glycerol. Final protein was aliquoted and stored at −80 °C.

### smFRET data analysis

The images were analyzed using MATLAB with an in-house program. Briefly, the fluorescence signals of Cy3 and Cy5 were averaged in a sliding window of 10 frames, and the fluorescence intensity of Cy3 was corrected for fluorescence leakage to calculate corresponding FRET values. The corrected Cy3 and Cy5 fluorescence intensities and FRET efficiency (E = I_acceptor_/(I_acceptor_+I_donor_) were plotted as a function of time. Molecules were selected that fit the following criteria: when the 638 nm laser was turned on at the end of the experiment, the Cy5 molecule was successfully excited, and Cy3 and Cy5 fluorescent signals showed anti-correlation. A FRET histogram was plotted using Origin 9 software. Gaussian fitting of FRET histograms was also performed using Origin 9. Histograms were fitted with a three-Gaussian model, first globally and then individually without constraints, to determine peak positions. Fractions of each FRET population were determined based on the area under each Gaussian. The dissociation constant (*K_d_*) was determined in GraphPad Prism 8 using the equation below: Y = B_max_*X*^h^*/(*K_d_^h^*+ X*^h^*), Bmax was less than 1, where X represents Cdc13 protein concentration, Y represents the bound fraction and *h* represents the hill coefficient. The average value and standard error of the mean were calculated from at least three independent experiments.

The time-tracing experiments recorded the images of DNA substrates in real-time after Cdc13 was added. The interval between protein addition and image focusing was defined as the dead time. In these experiments, the FRET efficiency values of each state were first determined based on the snapshot experiments. The time trajectory was then fitted using the vbFRET (3) program, which uses a Hidden Markov Model (HMM) and a variational Bayesian expectation maximization (VBEM) algorithm, yielding time series of discrete states. The fraction of transition events was calculated, and the transition rate calculations were described in the transition rate analysis.

### CoSMoS data analysis

The images were analyzed using Python with an in-house program adapted from Jeff Gelles’ lab (4). Briefly, DNA fluorescent signals from the first frame were averaged over the next 5-10 frames. The region encircling a distinct fluorescent spot was fitted to the spot center and designated as an area of interest (AOI). The AOIs from the DNA channel were then mapped onto the Cdc13 channel.

In snapshot experiments, colocalization events were defined as spots within 1.5 pixels of each other. The colocalization ratio was calculated by dividing the number of colocalized Cdc13-DNA spots by the total number of DNA spots and representing the bound fraction. This bound fraction was then applied to calculate the dissociation constant (*K_d_*). In the time-tracing experiment, mapped AOIs were used to measure changes in the fluorescence intensity over time. The interval between protein addition and the start of recording was defined as the deadtime. Time traces of fluorescence intensity were fitted with a Hidden Markov Model (HMM) by designing 3 states, producing a time series of discrete states. The fraction of transition events was calculated, and the transition rate calculations were described in the transition rate analysis. To calculate stoichiometry, intensity time traces were fitted using ebFRET program (5), which uses a Hidden Markov Model (HMM) with an Empirical Bayesian approach, allowing up to four maximum states to generate a time series of discrete states. The traces showing a stepwise decrease in intensity until complete bleaching were analyzed, with each state persisting for at least 2 frames.

### Transition rate analysis

With the state time series estimated by vbFRET, it is then possible to calculate the transition rates. We assume the reaction kinetics can be described by a 3-state continuous-time homogeneous Markov model, which can be defined by a transition rate matrix Q, with 6 parameters corresponding to the pseudo-first-order rate constants:

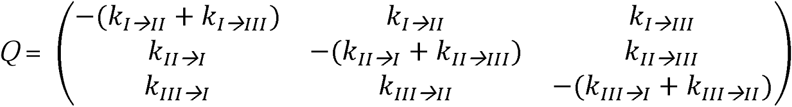

where ki→j is the transition rate from state i to state j if i ≠ j.

The goal of the analysis was to estimate the transition rate matrix Q from the state time series data. The R package msm ^6^ was then used to find the transition rate matrix that maximizes the likelihood of the state time series. Although the state at time 0 was not observed directly, we included the protein-free state at time 0 as the first observation because all the molecules must be protein-free before protein addition.

---

**Figure S1.**
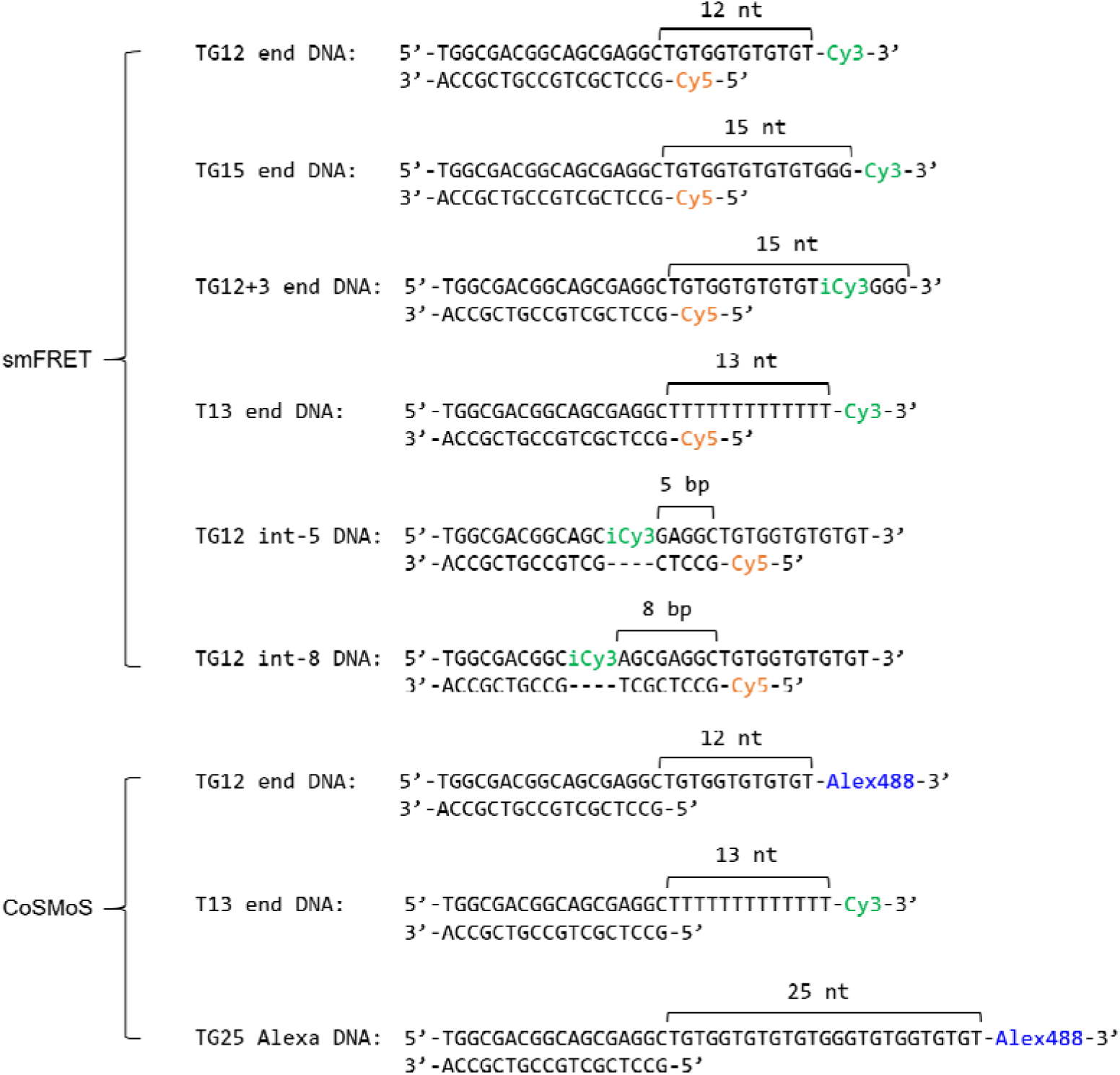
Sequences of DNA substrates used in this study.

**Figure S2.**
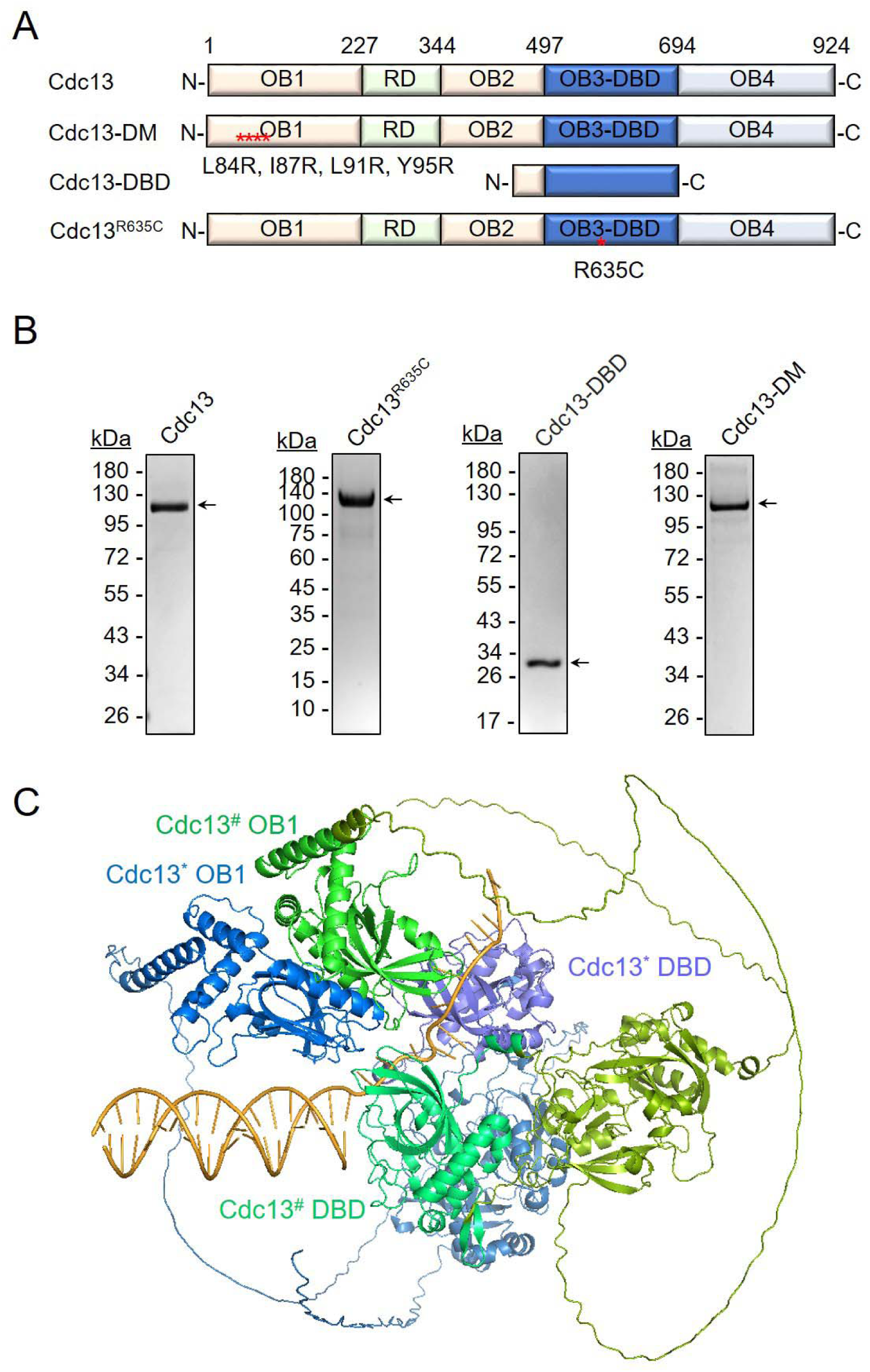
**(A)** Domain structure of Cdc13 and mutants. Cdc13-DM is the dimerization mutant, Cdc13-DBD contains only the DNA-binding domain, and Cdc13R635C is defective in DNA binding. **(B)** Purification of Cdc13 and its mutants. Wild-type Cdc13, Cdc13^R635C^, and Cdc13-DM with 6-His tag were purified from sf21. 6-His tagged Cdc13-DBD was isolated from *E. coli*. Coomassie Blue-stained 10% SDS-polyacrylamide gel of 1 µg each of purified Cdc13 is presented. **(C)** AlphaFold3-predicted model of a 2:1 Cdc13–TG12 complex. The two Cdc13 molecules are shown in shades of blue and green, respectively. The TG12-end DNA sequence is identical to that shown in Fig S1.

**Figure S3.**
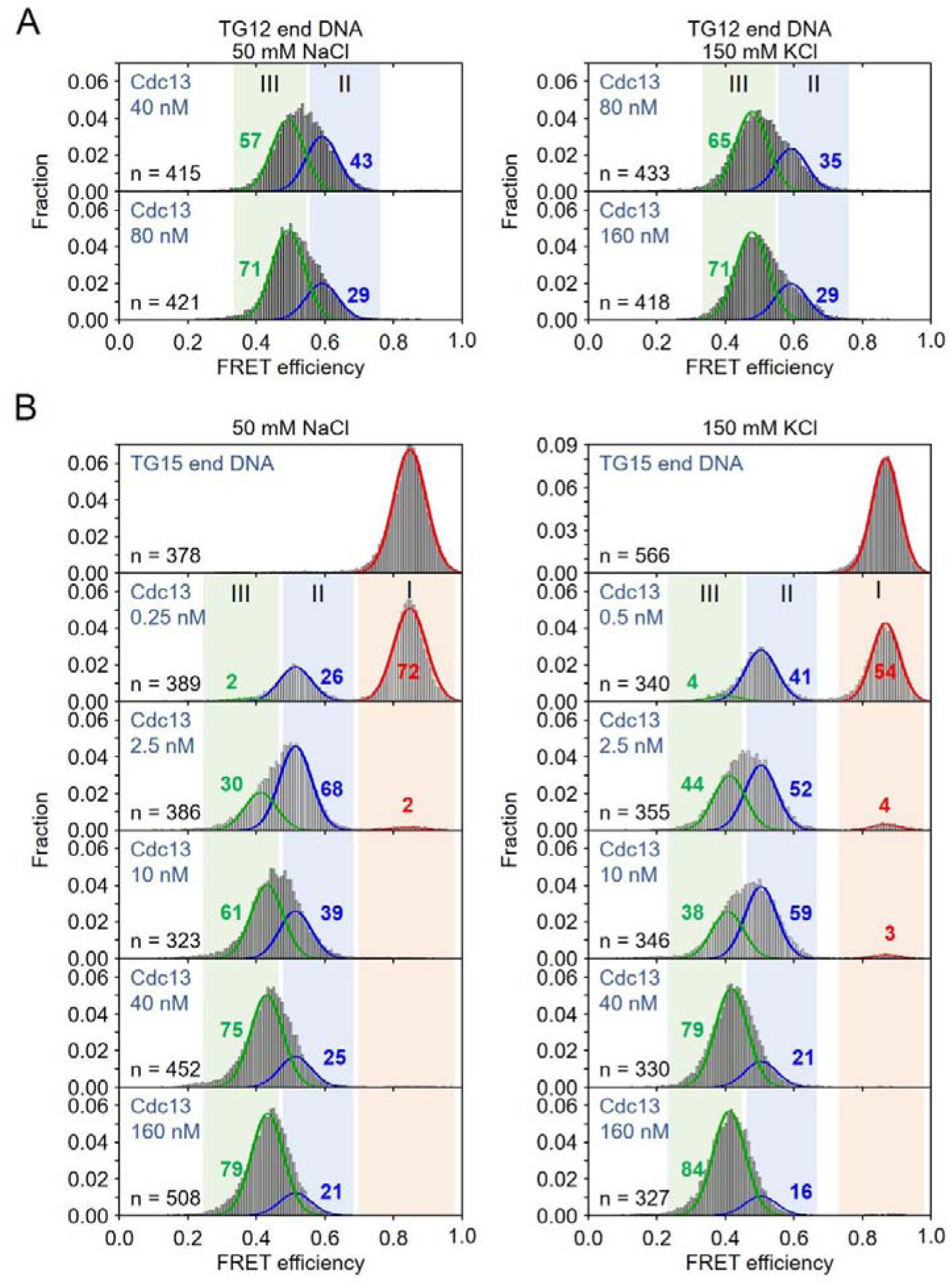
**(A)** FRET analysis of Cdc13 binding to TG12 end DNA. Histograms showing FRET efficiencies for TG12 end DNA with varying Cdc13 concentrations in 50 mM NaCl (left) and 150 mM KCl (right) separately. **(B)** FRET analysis of Cdc13 binding to TG15 end DNA. Histograms showing FRET efficiencies for TG12 end DNA with varying Cdc13 concentrations in 50 mM NaCl (left) and 150 mM KCl (right) separately. The fractions of state I (red), II (blue) and III (green) are shown across different Cdc13 concentrations.

**Figure S4.**
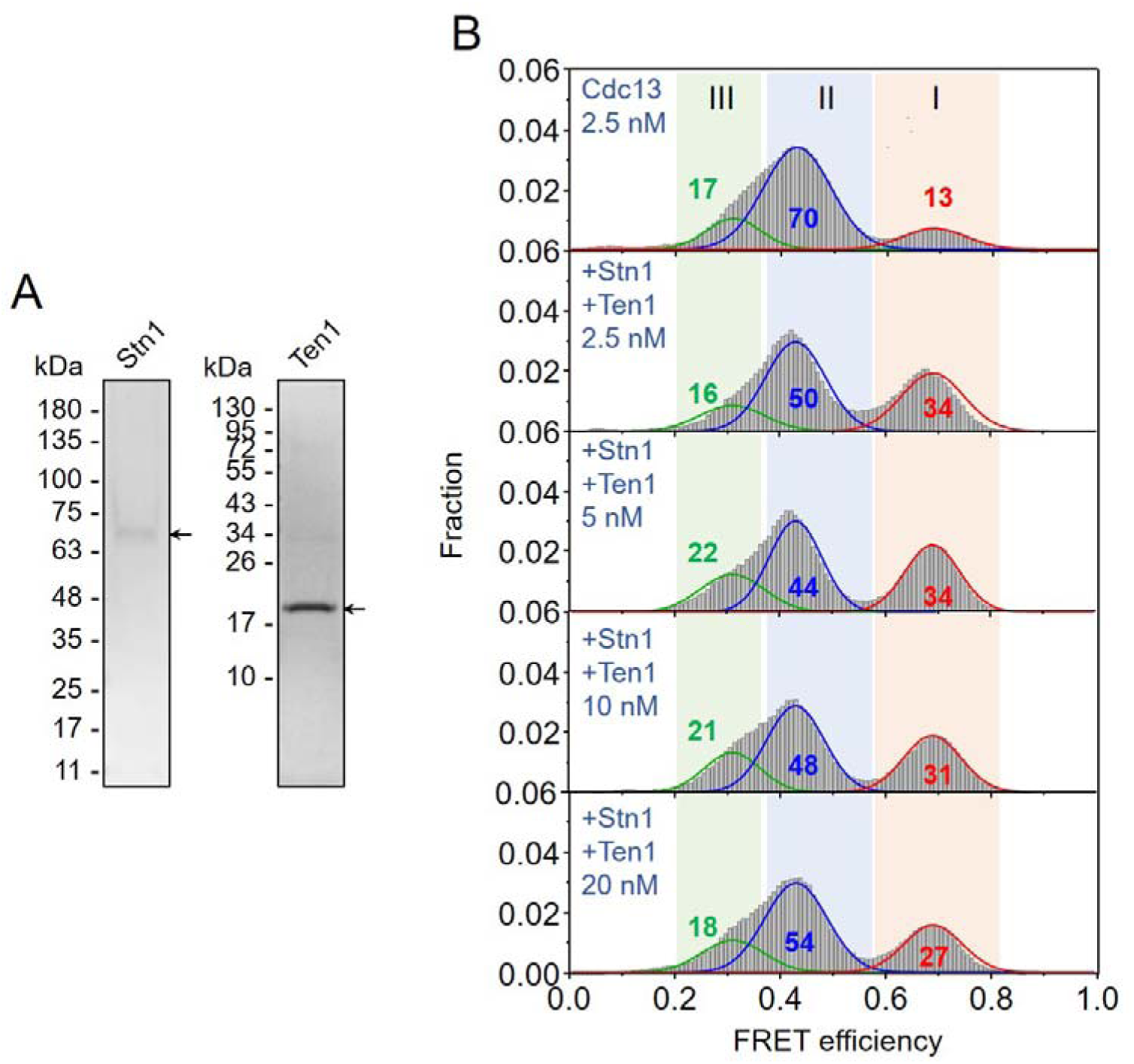
Formation of state II and III in the presence of Stn1 and Ten1. **(A)** Stn1 (TAP tagged) and Ten1 (6-His tagged) were purified from yeast cells and *E. coli*, respectively. Stn1 was analyzed by 8% SDS-PAGE, while Ten1 was analyzed by 12% SDS-PAGE. Coomassie Blue-stained gels showing the purified proteins are presented. **(B)** Stn1 and Ten1 did not appear to affect the formation of state II and III. FRET histograms of TG12-end DNA incubated with 2.5 nM Cdc13 alone or with the indicated concentrations of Stn1 and Ten1 are shown.

**Figure S5.**
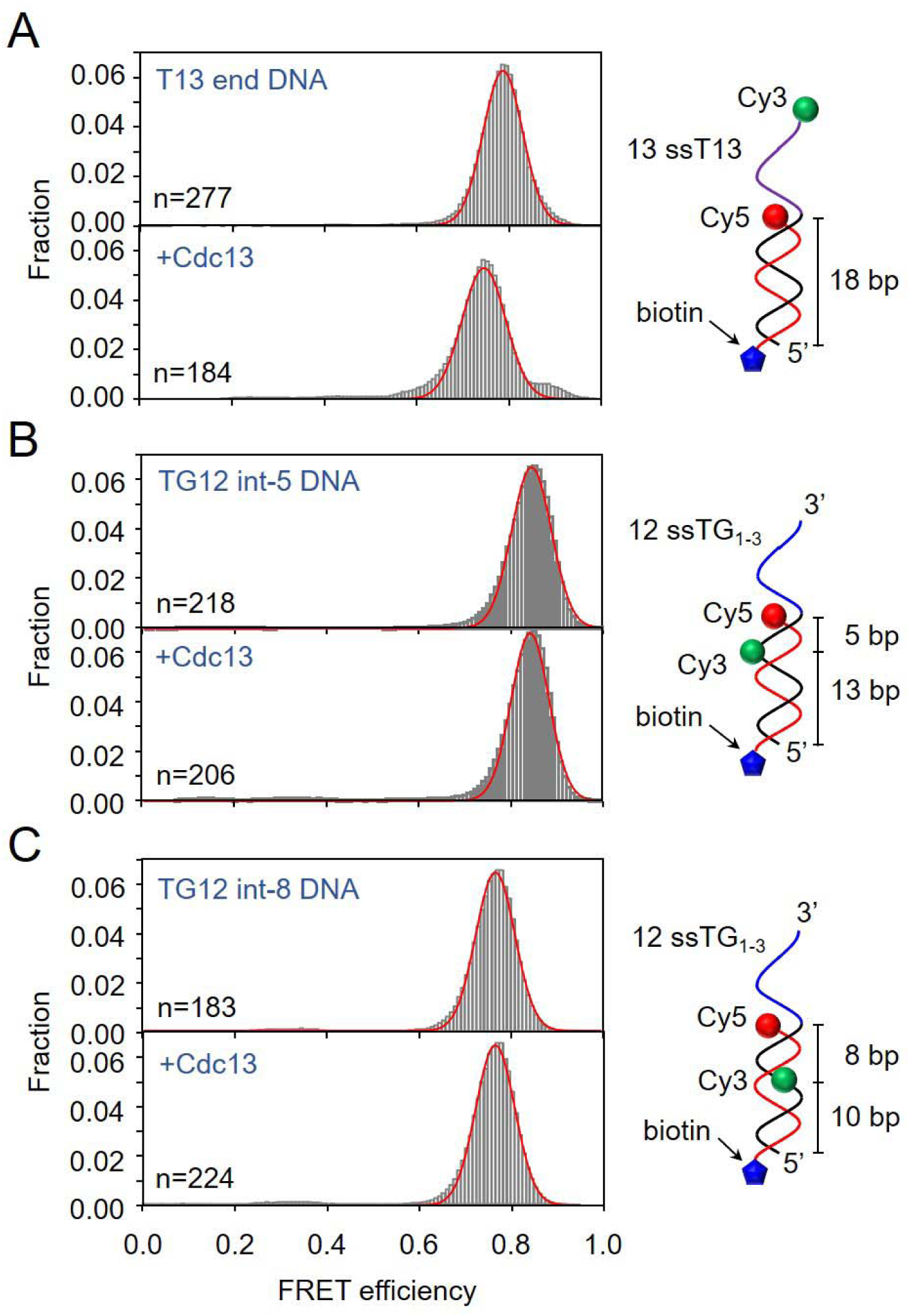
FRET analysis of Cdc13 binding to T13 end, TG12 int-5 DNA and TG12 int-8 DNA. Histograms of FRET efficiencies for **(A)** T13 end DNA, **(B)** TG12 int-5 DNA, and **(C)** TG12-int8 DNA in the absence of Cdc13 (top) or 10 nM Cdc13 (bottom). Schematics on the right show the design of DNA substrates. Cdc13 does not result in FRET change in these substrates.

**Figure S6.**
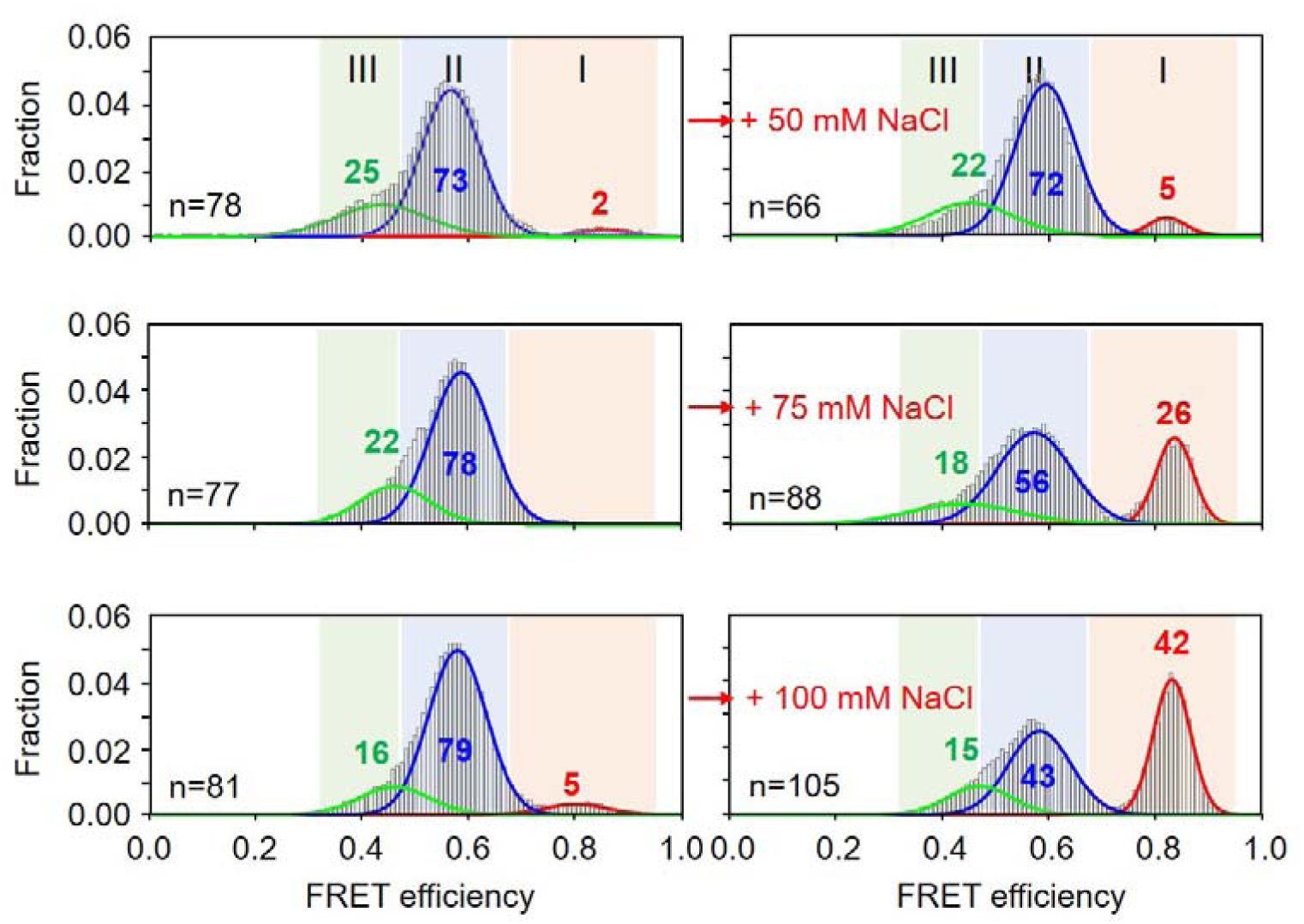
Exemplary FRET histograms of salt challenge experiments as analysis shown in Fig. 2A. High concentration of wild-type Cdc13 is loaded onto TG12-end DNA (histograms shown on the left), challenged with indicated NaCl concentrations (histograms shown on the right).

**Figure S7.**
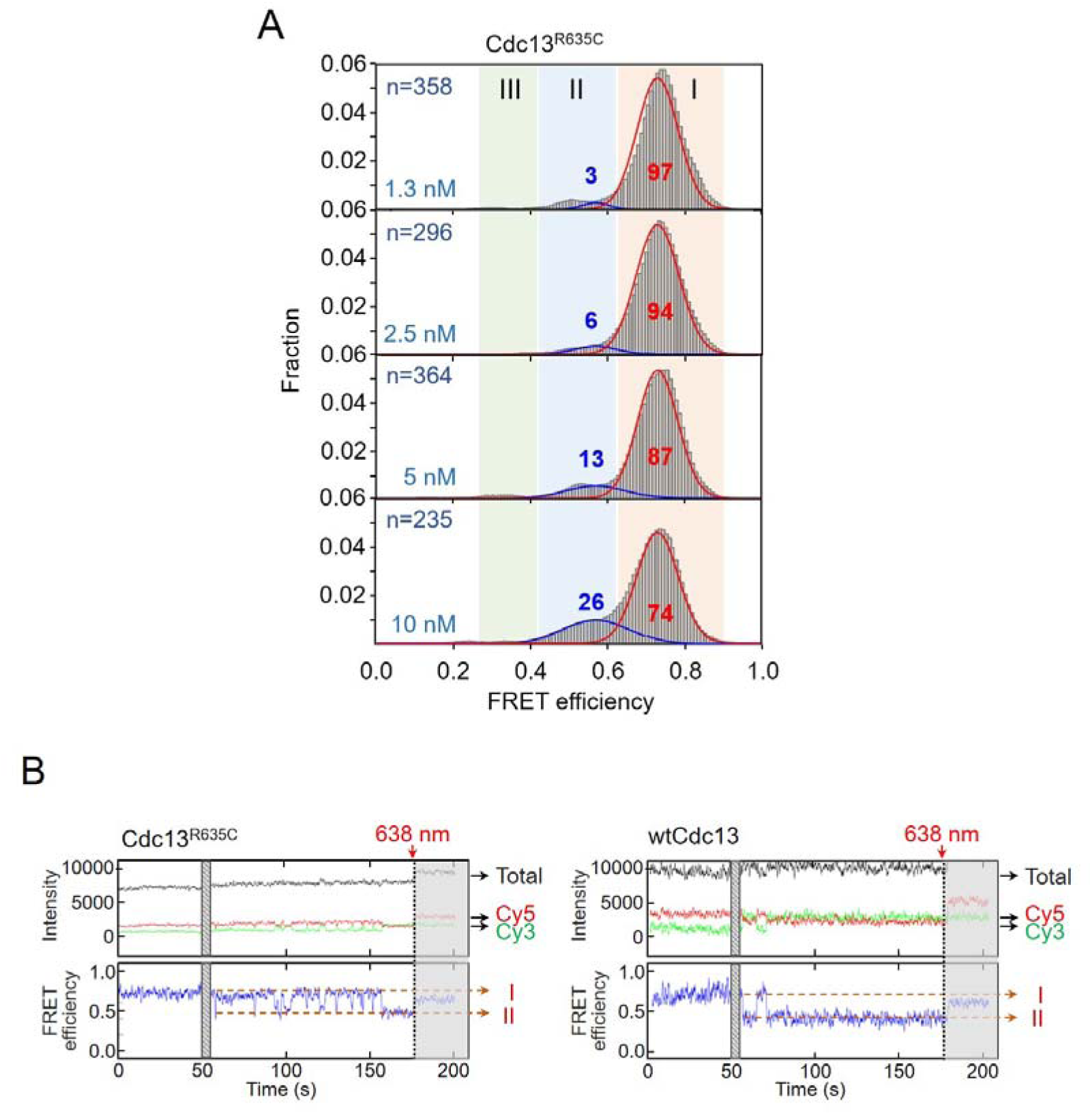
DNA-binding of Cdc13 is required for forming state II populations. **(A)** DNA binding-defective mutant of Cdc13^R635C^ severely reduces state II formation. FRET histograms of indicated amounts of Cdc13^R635C^ with TG12-end DNA are presented. **(B)** Representative smFRET trajectory showing state II formation by high concentration of Cdc13^R635C^ (left) is not stable compared to wtCdc13 (right). Total fluorescence, donor Cy3, and acceptor Cy5 intensities are colored in black, green, and red, respectively. Corresponding FRET efficiencies are shown below in blue.

**Figure S8.**
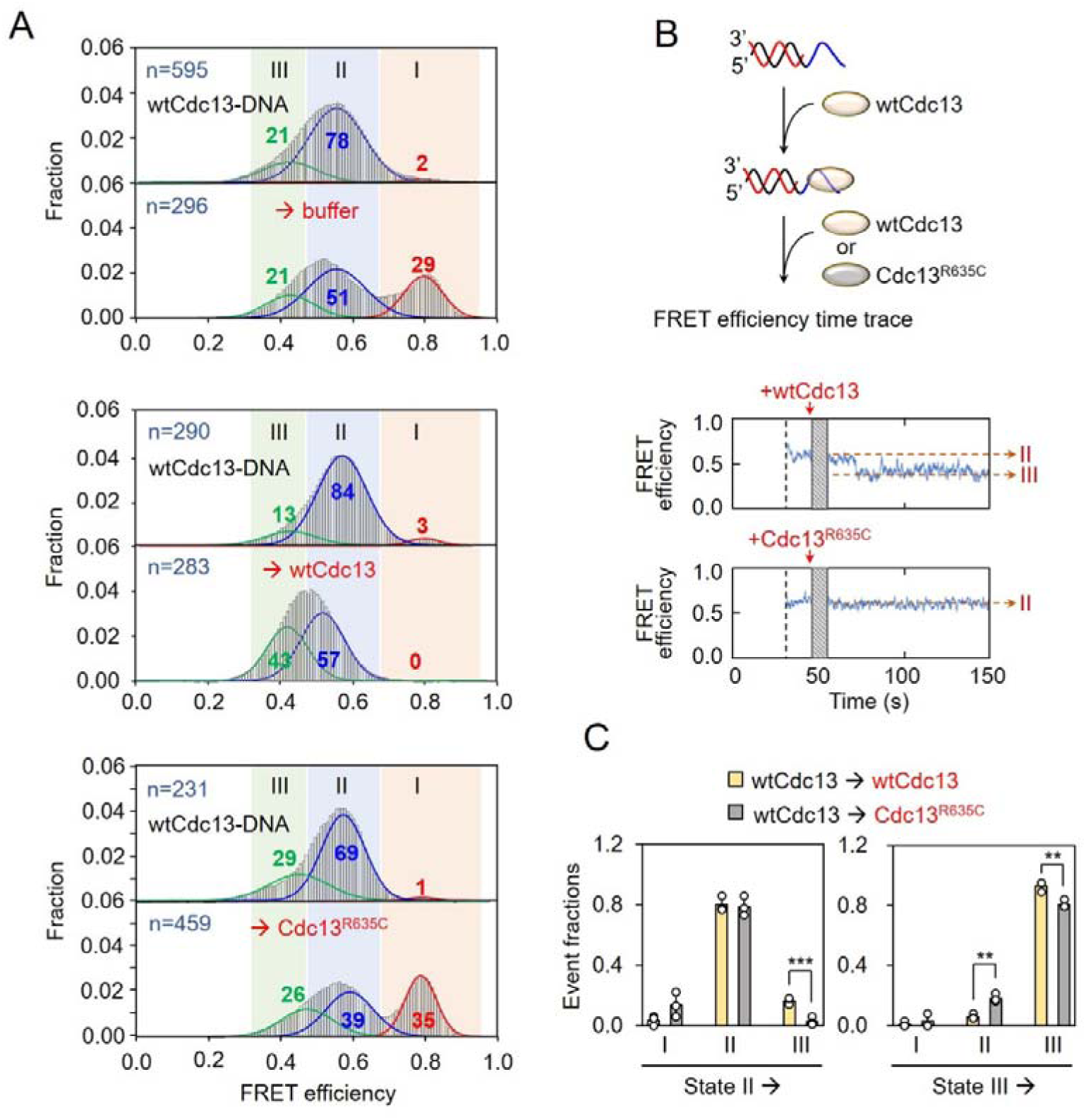
Details of Cdc13 chasing experiments shown in Fig. 2I. **(A)** Representative histograms of FRET analysis in Fig. 2I are presented. Cdc13 is loaded onto TG12 end DNA, then buffer-only control (top) or equal amount of either wtCdc13 (middle) or Cdc13^R635C^ mutant (bottom) is loaded. **(B)** Real-time analysis of Cdc13 challenging experiment and representative single-molecule FRET traces showing the transition from state II upon challenging with Cdc13^R635C^ (top) or wtCdc13 (bottom). FRET efficiencies are shown in blue. Dashed red lines correspond to different FRET states. **(C)** Analysis of the fraction change of state I, II, and III upon wtCdc13 or Cdc13^R635C^ mutant challenges. Upon Cdc13 challenging, more state II to III transitions are seen. An unpaired and two-tailed student’s t-test was applied to assess whether the means of the two groups were statistically different from each other, with ** denoting p<0.01.

**Figure S9.**
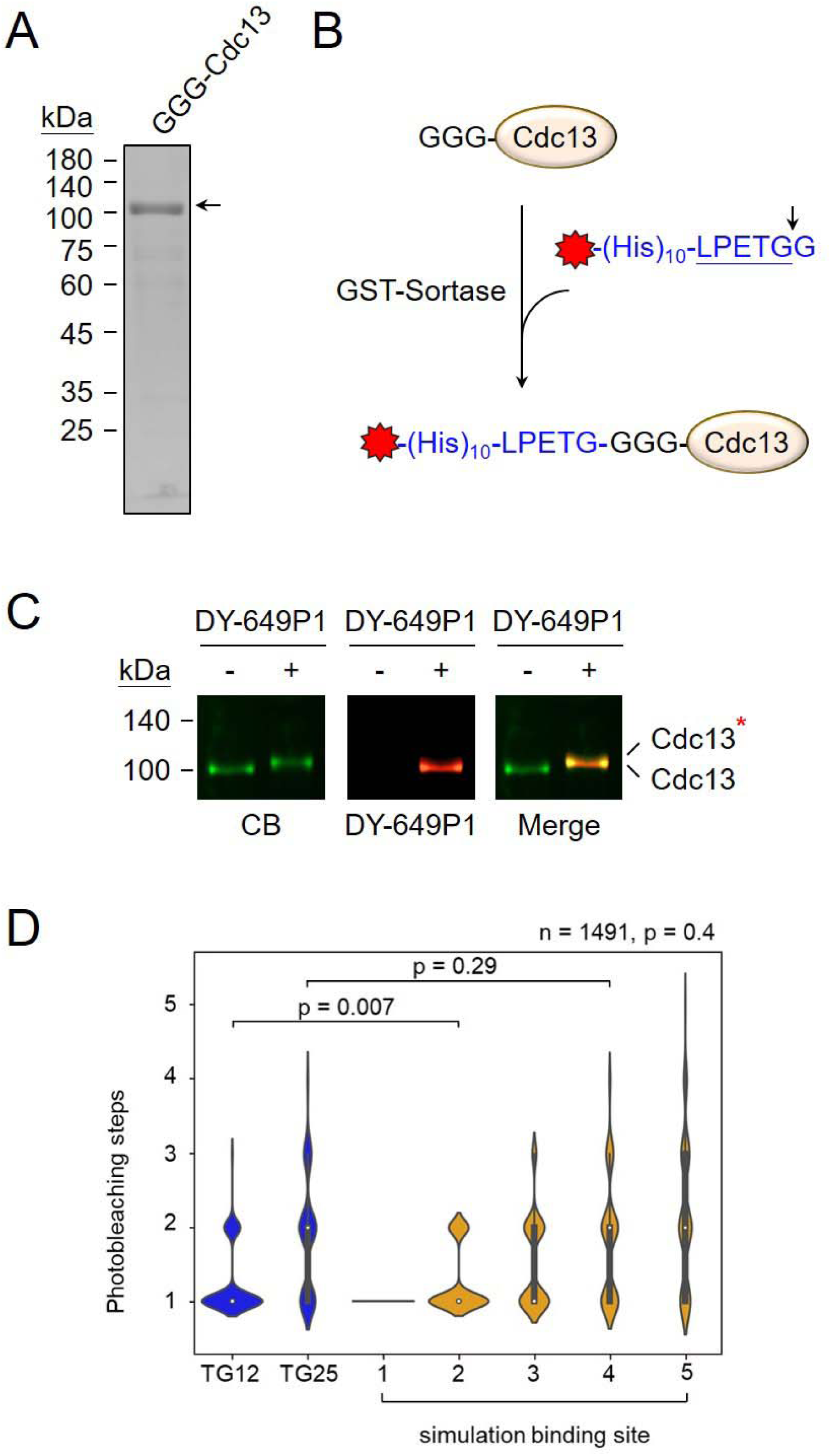
Purification and labeling of Cdc13. **(A)** 1 µg of isolated GGG-Cdc13 is analyzed by 10% SDS-PAGE. The Coomassie blue-stained gel is presented. **(B)** Schematic of the sortase-mediated Cdc13 labeling. **(C)** Fluorescent detection of DY649-labeled Cdc13. The gel stained by Coomassie blue (CB), DY-649, and merge images are shown. **(D)** Violin plots show the experimental results (blue) for TG12 and TG25 and the binomial distribution simulation results (yellow). Statistical significance was assessed using a chi-square goodness-of-fit test. Binomial random variables were generated using NumPy’s random.binomial function in Python, with the fluorescence labeling efficiency (p) as the probability of success and the number of Cdc13 molecules per DNA molecule (1–5) as the number of trials. Simulations were repeated until the number of simulated DNA molecules matched that of the TG12 experimental group (n).

**Figure S10.**
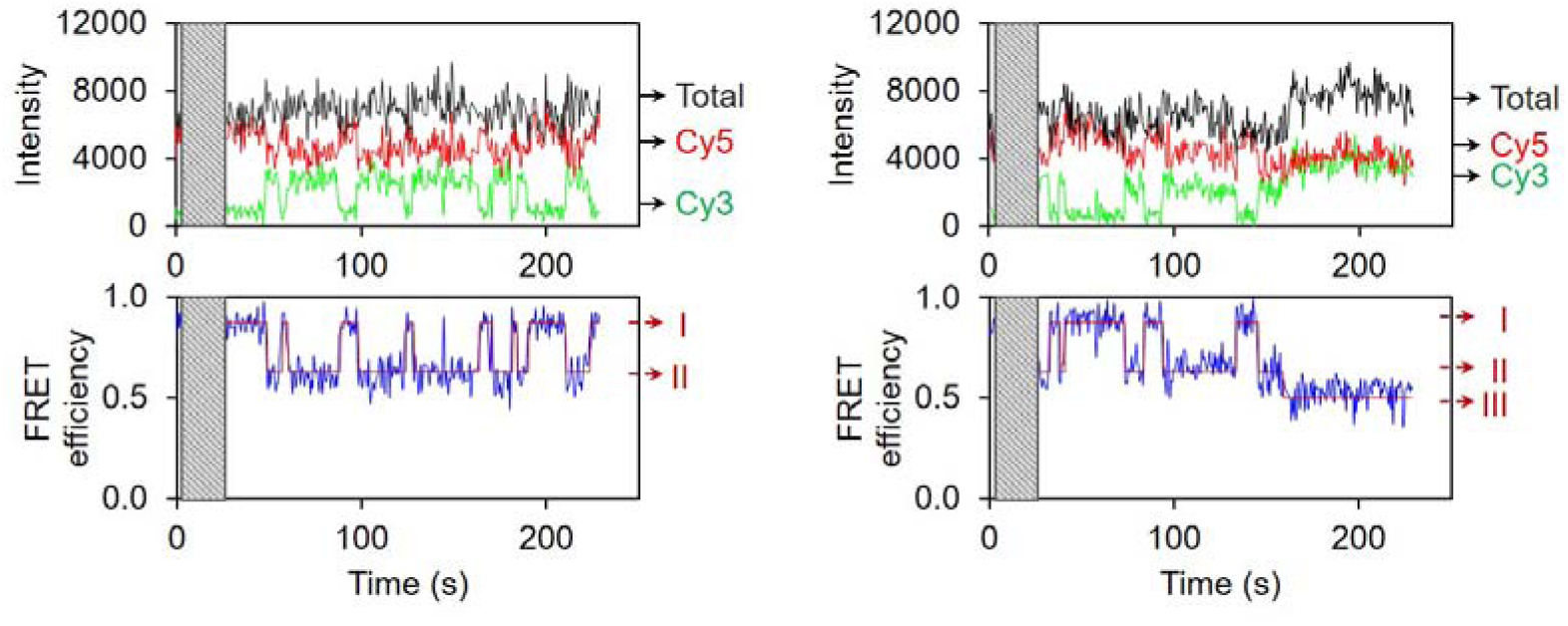
Representative single-molecule FRET time-course showing dynamic conversion among FRET states I, II, and III upon Cdc13 binding to TG12 end DNA in 150 mM KCl. Total fluorescence, donor Cy3, and acceptor Cy5 intensities are colored in black, green, and red, respectively. Corresponding FRET efficiencies are shown below in blue. Dashed red lines correspond to three FRET states. Grey block represents the unobserved dead time.

**Figure S11.**
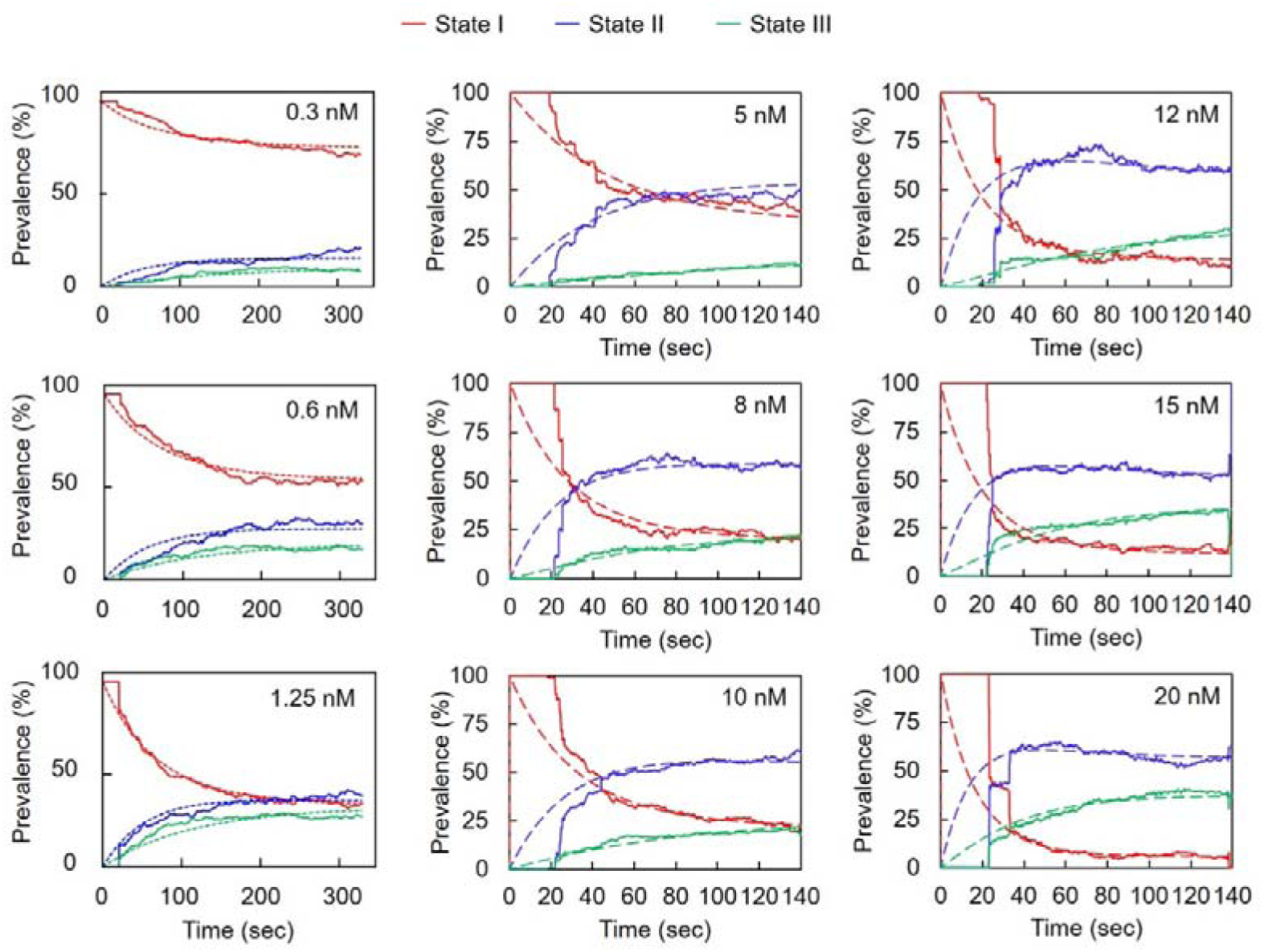
Transition rate analysis among various wtCdc13 binding states on TG12 end DNA in 50 mM NaCl based on the 3-state continuous-time homogeneous Markov Model. State time series were used to analyze the transition dwell times of different Cdc13 binding states. The Cdc13 concentrations are indicated in the figures. 0.3, 0.6, and 1.25 nM Cdc13 data were obtained from CoSMoS, and 5-20 nM data were obtained from FRET experiments. Prevalence of FRET state I (red, 0-Cdc13 bound), state II (blue, 1-Cdc13 bound), and state III (green, 2-Cdc13 bound) are presented (experimental data in solid lines and Markov model in dashed lines). At each Cdc13 concentration, 6 rates (I → II, II → III, I → III, II → I, III → II, and III → I) were determined, and four of them are shown in Fig. 4D.

**Figure S12.**
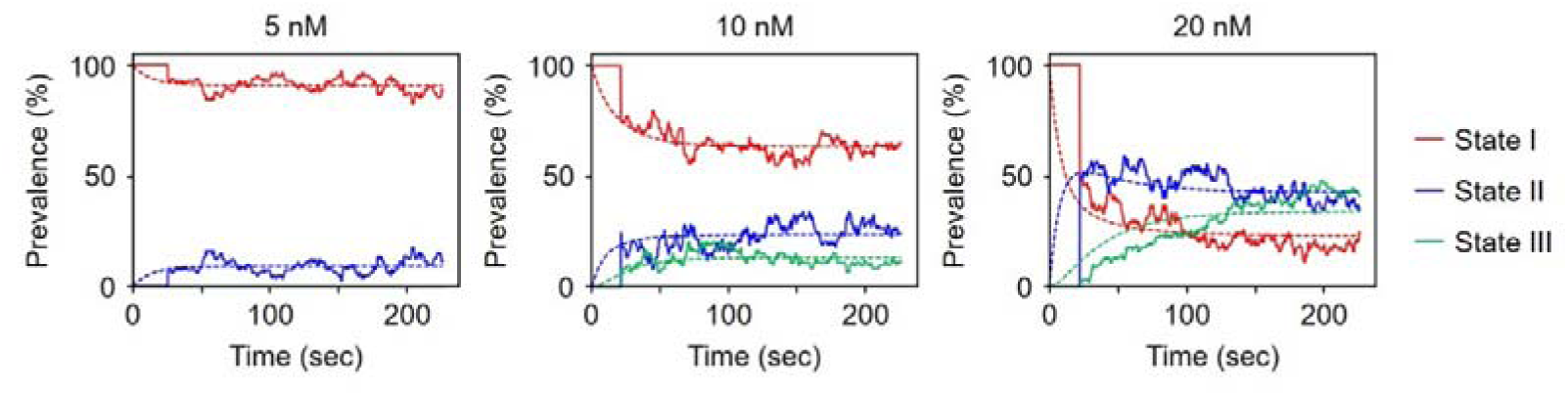
Transition rate analysis among various wtCdc13 binding states on TG12 end DNA in 150 mM KCl based on the 3-state continuous-time homogeneous Markov Model. State time series were used to analyze the transition dwell times of different Cdc13 binding states. Prevalence of FRET state I (red), state II (blue), and state III (green) are presented. At each Cdc13 concentration, 6 rates (I→II, II→III, I→III, II→I, III→II, and III→I) were determined, and two of them are shown in Fig. 4D (bottom).

**Figure S13.**
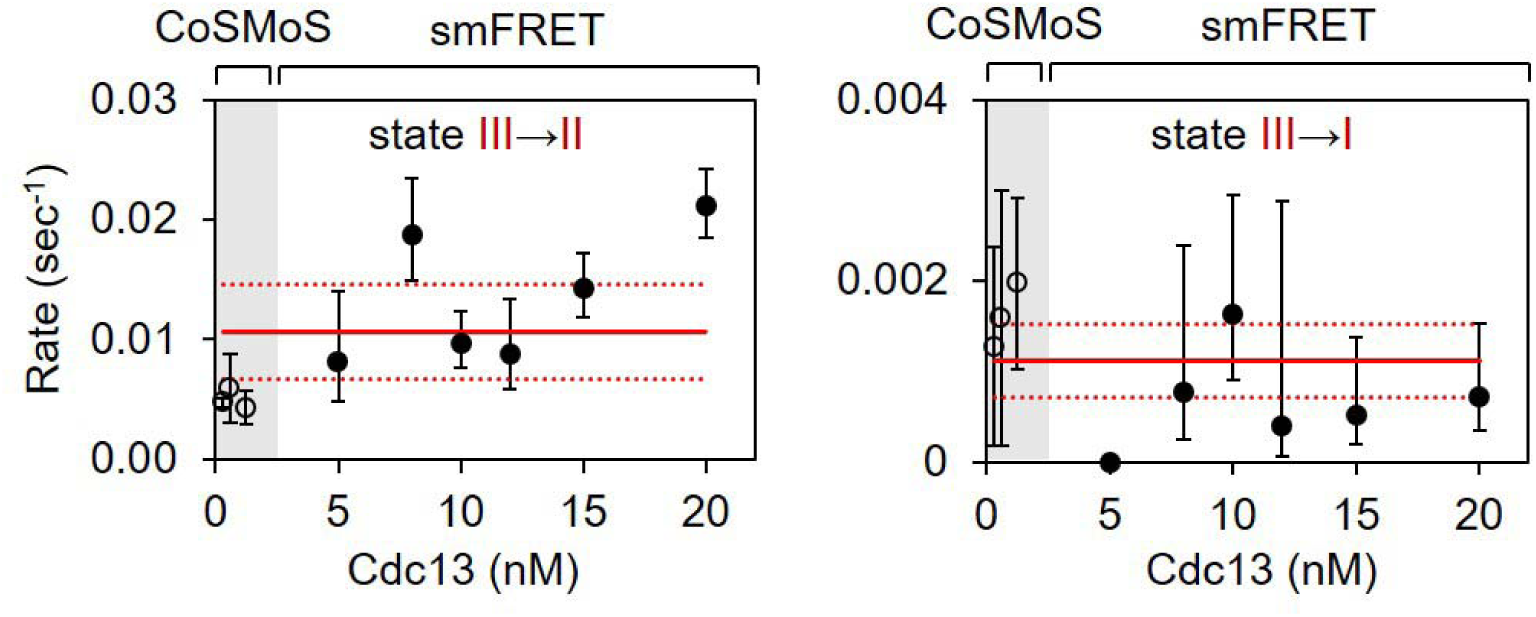
Transition rates (III→II, and III→I) were determined from Fig. S12, and its concentration dependence is shown. Note that the transition events exiting from state III (III→II, and III→I) are rare, as shown in Fig. 4C (middle), and thus large error bars are seen. Even though these rates are not as reliable as those presented in Fig. 4C due to the limited events observed, these rates showed weak concentration dependence, as expected. The 95% confidence intervals are shown as red dashed lines around the fitted trend line.

**Figure S14.**
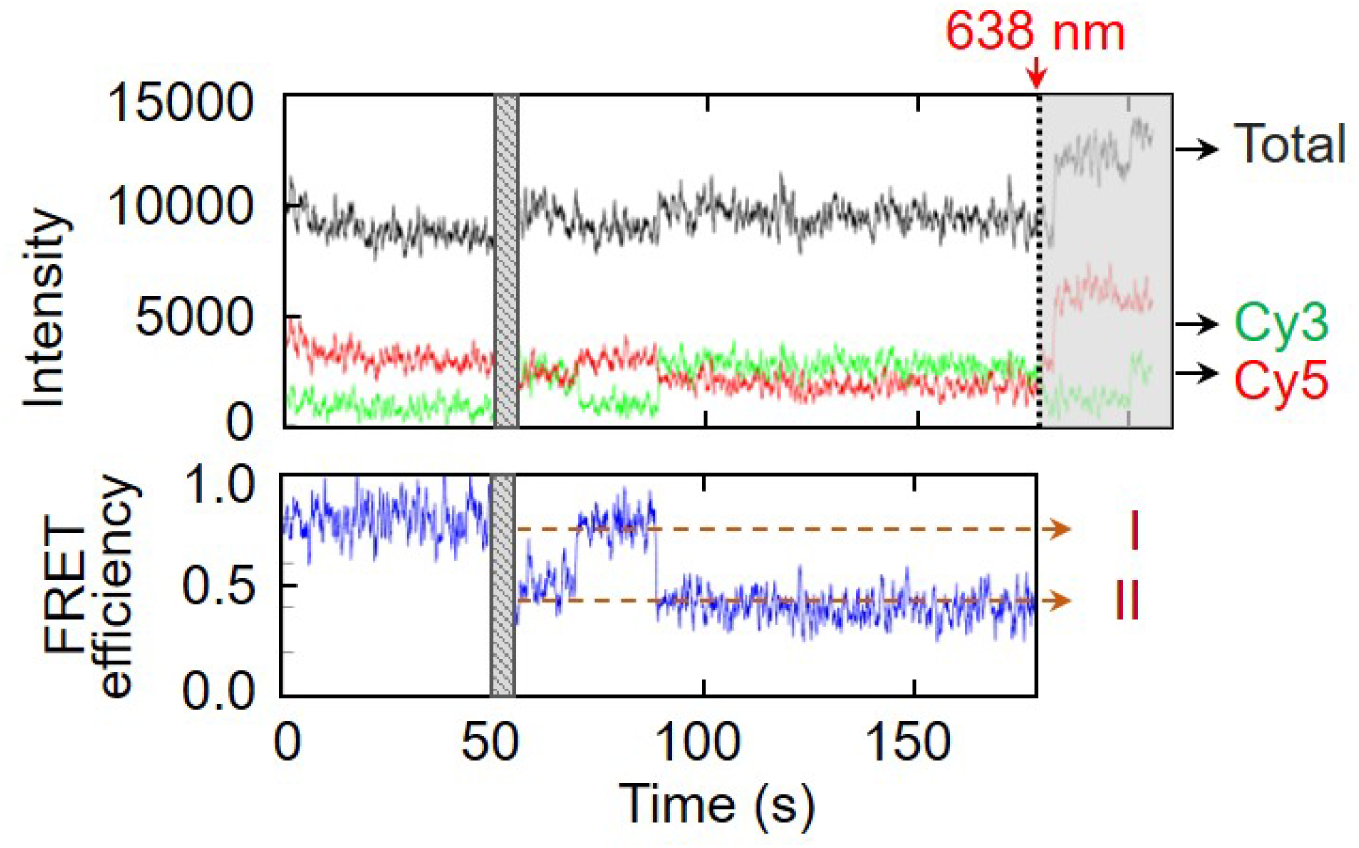
Transitions between Cdc13-DM binding states. Most transition alternations seen for dimerization-defective Cdc13-DM mutants are between state I and II.

**Figure S15.**
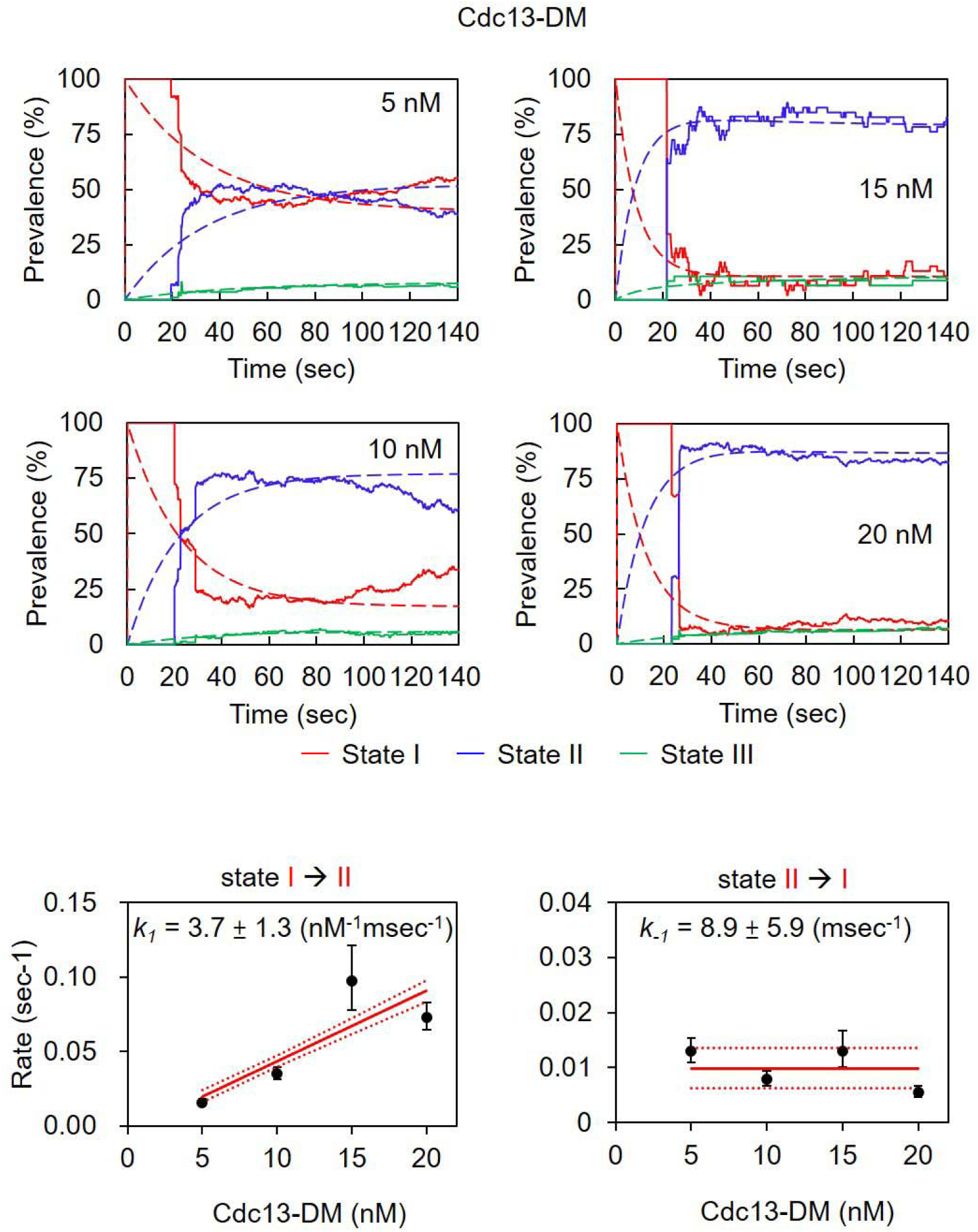
Transition rate analysis for Cdc13-DM mutants on TG12 end FRET DNA based on the 3-state continuous-time homogeneous Markov Model, as described in Fig. S12. Prevalence of FRET state I (red), state II (blue), and state III (green) are presented (experimental data in solid lines and Markov model in dashed lines). Transition rates (I→II and II→I) were determined, and their concentration dependence is shown.

**Table S1.** DNA molecule counts in Fig.4C-D.

| wtCdc13 | CoSMoS (50 mM NaCl) |  |  |  |  |  |  |
|  | r1 | r2 | r3 |  |  |  |  |
|---|---|---|---|---|---|---|---|
| 0.3 nM | 491 | 125 | 327 |  |  |  |  |
| 0.6 nM | 349 | 127 | 241 |  |  |  |  |
| 1.25 nM | 359 | 254 | 143 |  |  |  |  |
| wtCdc13 | FRET (50 mM NaCl) |  |  |  |  |  |  |
|  | r1 | r2 | r3 | r4 | r5 | r6 |  |
| 5 nM | 37 | 108 | 194 |  |  |  |  |
| 8 nM | 69 | 69 | 62 |  |  |  |  |
| 10 nM | 63 | 67 | 41 | 51 | 68 | 59 |  |
| 12 nM | 43 | 43 | 36 |  |  |  |  |
| 15 nM | 104 | 34 | 116 |  |  |  |  |
| 20 nM | 207 | 38 | 21 |  |  |  |  |
| wtCdc13 | FRET (150 mM KCl) |  |  |  |  |  |  |
|  | r1 | r2 | r3 | r4 | r5 | r6 | r7 |
| 5 nM | 83 | 101 | 71 | 93 | 50 | 66 | 82 |
| 10 nM | 86 | 105 | 117 | 120 | 104 | 51 | 74 |
| 20 nM | 143 | 89 | 97 | 90 | 72 | 44 | 90 |

**Table S2.** Summary of dissociation constants for TG12-end DNA binding.

| condition | Cdc13 | apparent *K<sub>d</sub>* (nM) | *k<sub>1</sub>* (nM<sup>-1</sup> msec<sup>-1</sup>) | *k<sub>-1</sub>* (msec<sup>-1</sup>) | kinetically derived *K<sub>d</sub>* (nM) |
|---|---|---|---|---|---|
| 50 mM NaCl | WT | 1.2 | 2.5 | 9.2 | 3.7 |
|  | DM | 3.1 | 3.7 | 8.9 | 2.4 |

## Notes

### Competing Interest Statement

The authors have declared no competing interest.

### Summary of Updates

Figure 2 was reorganized. All revisions were made in response to the comments and suggestions from both reviewers.

